# Protectin DX resolves fracture-induced postoperative pain in mice via neuronal signaling and GPR37-activated macrophage efferocytosis

**DOI:** 10.1101/2025.11.10.687700

**Authors:** Yize Li, Sangsu Bang, Jasmine Ji, Jing Xu, Min Lee, Sharat Chandra, Charles N Serhan, Ru-Rong Ji

## Abstract

Protectin DX (PDX) is a member of the superfamily of specialized pro-resolving mediators (SPMs) and exerts anti-inflammatory actions in animal models; but the specific signaling pathways involved in its actions have not been fully elucidated. Here we demonstrate the analgesic actions of PDX in a mouse model of tibial fracture-induced postoperative pain (fPOP) via activation of GPR37. Intravenous early-phase and late-phase treatment of PDX (100 ng/mouse) effectively alleviated fPOP. Compared to protectin D1/neuroprotectin D1, DHA, steroid, and meloxicam, PDX produced superior pain relief. While dexamethasone and meloxicam prolonged fPOP, PDX shortened the pain duration. The analgesic effects of PDX were abrogated in *Gpr37*^−/−^ mice, which displayed deficits in fPOP resolution. PDX binds GPR37 and induces calcium responses in peritoneal macrophages. LC-MS-MS-based lipidomic analysis revealed that endogenous PDX levels were ∼10-fold higher than PD1 in muscle at the fracture site. PDX promotes macrophage polarization via GPR37-dependent phagocytosis and efferocytosis through calcium signaling *in vitro*, and it further enhances macrophage viability and efferocytosis *in vivo* via GPR37. Finally, PDX rapidly modulates nociceptor neuron responses by suppressing C-fiber-induced muscle reflex *in vivo* and calcium responses in DRG neurons *ex vivo* and reducing TRPA1/TRPV1-induced acute pain and neurogenic inflammation *in vivo*. Our findings highlight multiple benefits of PDX to manage postoperative pain and promote perioperative recovery.

## Introduction

Fracture-related postoperative pain (fPOP) is associated with tissue injury in joints and muscles, local inflammation, as well as nerve injury and neuroinflammation in the peripheral and central nervous systems (1, 2). Although multimodal analgesia is widely used, postoperative pain after orthopedic surgery is still frequently undertreated, hindering recovery (3). Opioids have traditionally been the mainstay for postoperative pain management. However, concerns over their side effects, such as the potential for abuse and respiratory suppression, which drive the ongoing opioid epidemic (4), have created an urgent need to optimize non-opioid analgesic options (3, 5). Furthermore, anti-inflammatory treatments such as steroids and non-steroidal anti-inflammatory drugs (NSAIDs) produce gastrointestinal complications and may further impair the resolution of inflammatory pain and low-back pain (6, 7). Thus, there is an urgent need to develop more effective and safe pain therapeutics.

The pro-resolution pharmacology was proposed for controlling inflammation and pain (8), based on the potent analgesic actions of resolvins (1-300 ng/mouse) via different administration routes, in animal models of pain (9). Specialized pro-resolving mediators (SPMs) consist of omega-3 eicosapentaenoic acid (EPA)-derived E-series resolvins, docosahexaenoic acid (DHA)-derived D-series resolvins, protectins and maresins, cysteinyl-SPMs, as well as omega-6 arachidonic acid-derived lipoxins (10). DHA is crucial for brain development (11), and DHA-derived SPMs consist of resolvin D1-D6 (RvD1-RvD6), maresin 1-2 (MaR1-MaR2), and protectin D1 (PD1) (12, 13). Given its critical role in neuroprotection, PD1 is also termed neuroprotectin D1 (PD1/NPD1) (14). SPMs, produced during the resolution phase of inflammation, bind to specific G protein-coupled receptors (GPCRs) and regulate the activities of different cell types, including neutrophils, macrophages, T cells, epithelial cells, and neurons, promoting inflammation resolution, tissue repair, and pain relief (15, 16) (Supplementary Figure 1A). Mechanistically, SPMs resolve inflammation via macrophage phagocytosis and efferocytosis (17). PD1/NPD1 was shown to resolve inflammatory pain and infection-induced pain through GPR37-mediated macrophage phagocytosis (18, 19). Notably, pain resolution or chronicity is associated with SPM production (20, 21).

Protectin DX (PDX) was identified as a stereoisomer of PD1 in 2006 (22), and was further chemically characterized in 2009 (23). Compared to PD1, PDX has been shown to exhibit different chemical and biological properties (24). For example, PDX (10S,17S-diHDHA) is biosynthesized via double dioxygenation determined using molecular oxygen (^18^O) incorporation in PDX and mass spectrometry, and was shown to reduce neutrophil infiltration in mouse peritonitis *in vivo*, albeit less potent than PD1 (22) (Supplementary Figure 1B). PDX alleviates insulin resistance and Type II diabetes (25) and inhibits blood platelet activity at higher concentrations (μM range) (23, 26). However, key questions still remain regarding the therapeutic potential of PDX in pain management: the receptor for PDX is unknown; endogenous PD1 and PDX levels in health and disease have not been characterized; optimal pain models, delivery routes, and dosing for PDX remain undefined, with SPM analgesic efficacy varying by models (20, 27) and sex (28); and the immune-neuronal mechanisms of PDX are largely unexplored. To this end, we tested the inhibitory actions of PDX on the development, maintenance, and recovery of fPOP and further investigated the molecular and cellular mechanisms using a clinically relevant orthopedic surgery pain model (29). Our findings demonstrate that intravenously administered PDX, at low doses (30-100 ng/mouse), can potently mitigate fPOP and promote the resolution of postoperative via GPR37.

## Results

### Early perisurgical treatment of PDX mitigates the development of fracture-induced postoperative pain (fPOP)

PDX and PD1/NPD1 were obtained from Cayman Chemical and validated using UV spectroscopy and mass spectrometry, matching the UV spectrums, retention times, and fragmentation patterns of the compounds to those of known standards (Supplemental Figures 2 and 3). Before testing the drug effects, we first investigated the time course of postoperative pain in male and female CD1 mice after sham surgery and tibial fracture surgery. Mice with sham surgery exhibited no changes in mechanical pain (paw withdrawal threshold, PWT, in von Frey tests, Supplemental Figure 4A), cold pain (acetone test, Supplemental Figure 4B), and heat pain (Hargreaves tests, Supplemental Figure 4C). However, following tibial fracture surgery, animals exhibited increased mechanical, cold, and heat pain on the first day, and these evoked pains persisted for 14 days and fully recovered by 21 days post-surgery (Supplemental Figure 4A-C). We also examined spontaneous pain by assessing guarding behavior, which appeared on the first day post-surgery, maintained on day 7, but returned to baseline on day 10 (Supplemental Figure 4D). The grimace test only showed spontaneous pain on days 3 and 5 post-fracture (Supplemental Figure 4E).

We next investigated the effects of PDX early-treatment on fPOP, administered during the peri-surgical period with the first injection given right before surgery and the second injection given 2 days after surgery (Figure 1A). We performed an intravenous dose-response study (1-100 ng per mouse), a standard peri-operative route (30). Von Frey testing showed no effects on PWT at 1-10 ng, a transient effect at 30 ng, and robust anti-nociception lasting for 5 hours at 100 ng (Supplemental Figure 5A); thus, 100 ng dose was used for most experiments in this study. Compared with PD1/NPD1, 300 ng NPD1 was required to achieve a similar effect to 100 ng PDX (Supplemental Figures 5A and 5B). ED_50_ analysis (52 ng vs. 205 ng) confirmed ∼4-fold greater potency of PDX over NPD1 (Supplemental Figure 5C).

**Figure 1.**
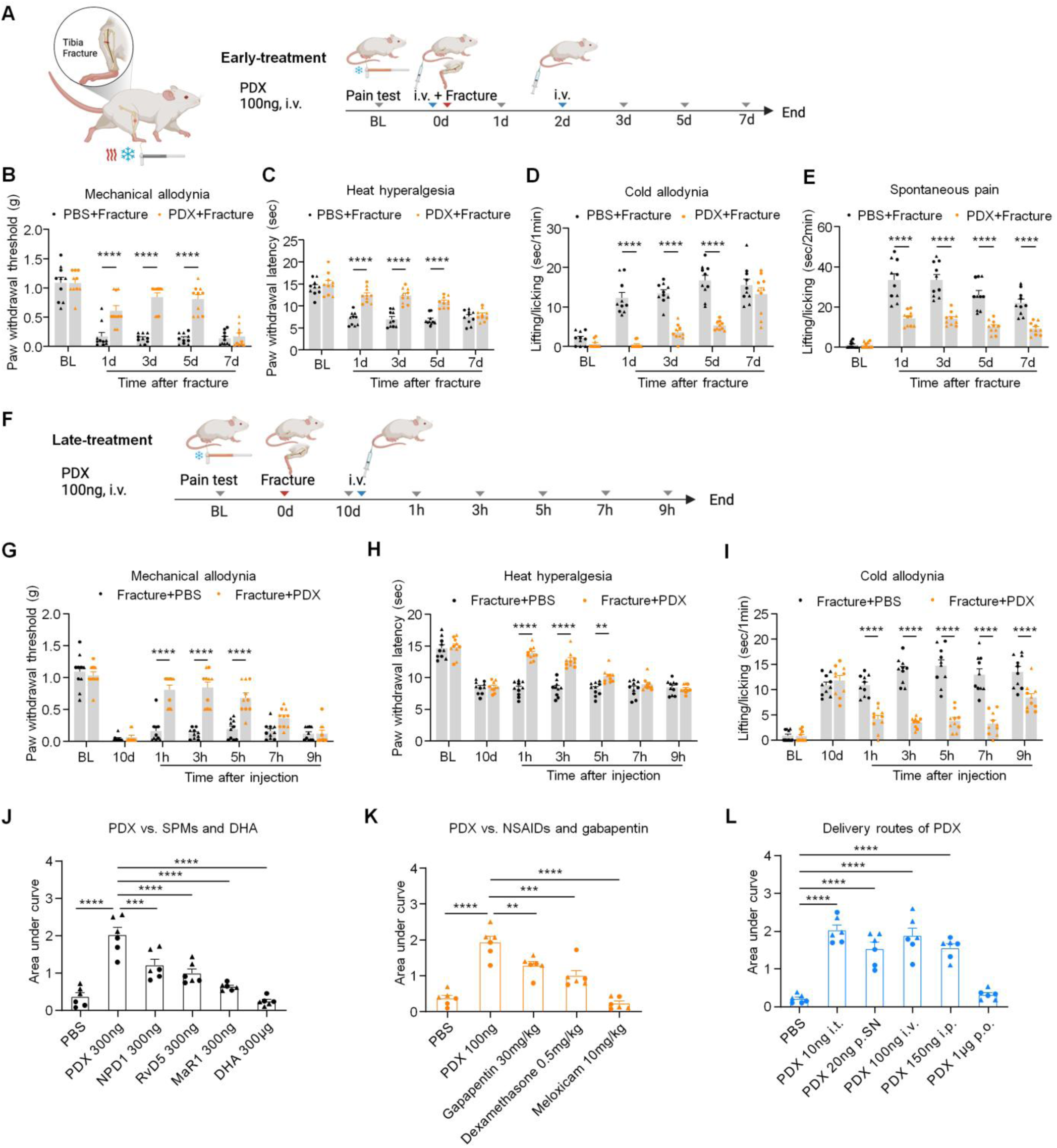
Perisurgical pre- and post-treatment with PDX attenuates tibial fracture-induced postoperative pain (fPOP) in CD1 mice. **(A**) Schematic of the tibial fracture model and PDX pre-treatment timeline. CD1 mice received two intravenous (i.v.) injections of PDX (100 ng, 100 μl) or PBS (100 μl): one immediately before fracture surgery and one post-fracture day 2. (**B-E**) Effects of PDX and vehicle on fPOP measured on post-surgical day 1, 3, 5, and 7, including mechanical pain (PWT in von Frey test, **B**), thermal pain (paw withdrawal latency in Hargreaves’ test, **C**), cold pain (duration of pain with lifting/licking behavior in acetone test, **D**), and spontaneous pain (duration of lifting/licking behavior, **E**). (**F**) Post-treatment paradigm. In a separate cohort, mice received a single i.v. injection of PDX (100 ng, 100 μl) or PBS on post-fracture day 10. (**G–I**) Post-treatment with PDX also reduced mechanical (G), thermal (H), and cold (I) pain. (**J-K**) Comparison of PDX with other SPMs (NPD1, RvD5, and MaR1; 300 ng/mouse) and DHA (300 μg/mouse, **J**), as well as anti-inflammatory drugs (dexamethasone, 0.5 mg/kg; meloxicam, 10 mg/kg) and gabapentin (30 mg/kg, **K**). (**L**) Route-of-administration comparison showing effective analgesia following intrathecal (i.t., 10 ng), peri-sciatic (p.SN, 20 ng), intravenous (i.v., 100 ng), intraperitoneal (i.p., 150 ng), or oral (p.o., 1 μg) delivery of PDX. Data in J-L are presented as Area under curve (AUC) of PWT. All data are represented as mean ± SEM. Statistics: two-way ANOVA with Bonferroni’s post hoc test (B-E, G-I) and one-way ANOVA with Tukey’s post hoc test (J-L). \*\**P* < 0.001, \*\*\**P* < 0.001, \*\*\*\**P* < 0.0001; *n* = 10 (B-E, G-I), *n* = 6 (J-L), and equal number of males and females are included; ▴ male, ● female.

We next evaluated early PDX treatment in fPOP during the first postoperative week across mechanical, thermal, cold, and spontaneous pain (Figure 1A). Intravenous PDX (100 ng) markedly reduced mechanical allodynia, thermal hyperalgesia, cold allodynia, and spontaneous pain on days 1, 3, and 5, and continued to suppress spontaneous pain on day 7 (all *P* < 0.0001; Figure 1B–E), with no sex differences (Supplemental Figure 5D). Thus, early i.v. PDX effectively attenuates acute fPOP.

### Late PDX treatment alleviates persistent fPOP and outperforms standard analgesics

Because fPOP persists for 2–3 weeks, we tested whether delayed PDX administration (100 ng, i.v., day 10) reduces established pain in male and female CD1 mice (Figure 1F). PDX significantly suppressed mechanical, thermal, and cold hypersensitivity at 1–5 h (all *P* < 0.0001 to *P* < 0.01) and produced prolonged relief of cold pain up to 9 h (Figure 1G–I), with no sex differences (Supplemental Fig. 5E). We next compared PDX with DHA, MaR1, NPD1, and RvD5 (all at 100 pmol, i.v.). PDX produced the greatest analgesic effect, with a significantly larger AUC than all comparators (*P* < 0.001; Figure 1J; Supplemental Fig. 6A). Post-operative pain is typically treated using dexamethasone (steroid), meloxicam (NSAID), and gabapentin (31). Notably, PDX (100 ng; ∼0.003 mg/kg) also outperformed gabapentin (30 mg/kg, i.p.) and dexamethasone (0.5 mg/kg, i.v.), despite 100–5000-fold lower doses; meloxicam (10 mg/kg, i.v.) was ineffective in late-phase fPOP (Figure 1K; Supplemental Figure 6B). Finally, PDX showed robust efficacy via intrathecal (10 ng), peri-sciatic (20 ng), intraperitoneal (150 ng), and i.v. delivery, each producing ∼5 h analgesia (Figure 1L; Supplemental Fig. 7). Oral gavage, even at 1000 ng, yielded only transient benefit. Together, these data demonstrate that delayed PDX treatment effectively reverses established fPOP, exhibits superior potency to other SPMs and clinical analgesics, and is effective across multiple delivery routes.

### PDX and anti-inflammatory treatments differentially regulate the duration of fPOP

Anti-inflammatory drugs such as steroids and NSAIDs may delay inflammatory pain resolution (6, 8). To compare their effects with PDX, we administered PDX (100 ng/mouse), dexamethasone (0.5 mg/kg), or meloxicam (5 mg/kg) immediately before surgery (Figure 2A). Dexamethasone produced transient analgesia (1–5 h), meloxicam showed delayed but sustained effects (3–24 h), and PDX elicited both rapid and lasting relief (1–24 h, *P* < 0.0001). AUC analysis indicated stronger acute antinociception with PDX and meloxicam than dexamethasone (Figure 2B). To assess longer-term benefit, a second dose was given at 48 hours (Figure 2C). Dexamethasone provided no sustained benefit and worsened pain at day 21; meloxicam produced transient relief but later increased pain (days 14–21). In contrast, PDX significantly improved pain from days 3–17 (*P* < 0.001), confirmed by AUC analysis (Figure 2D).

**Figure 2.**
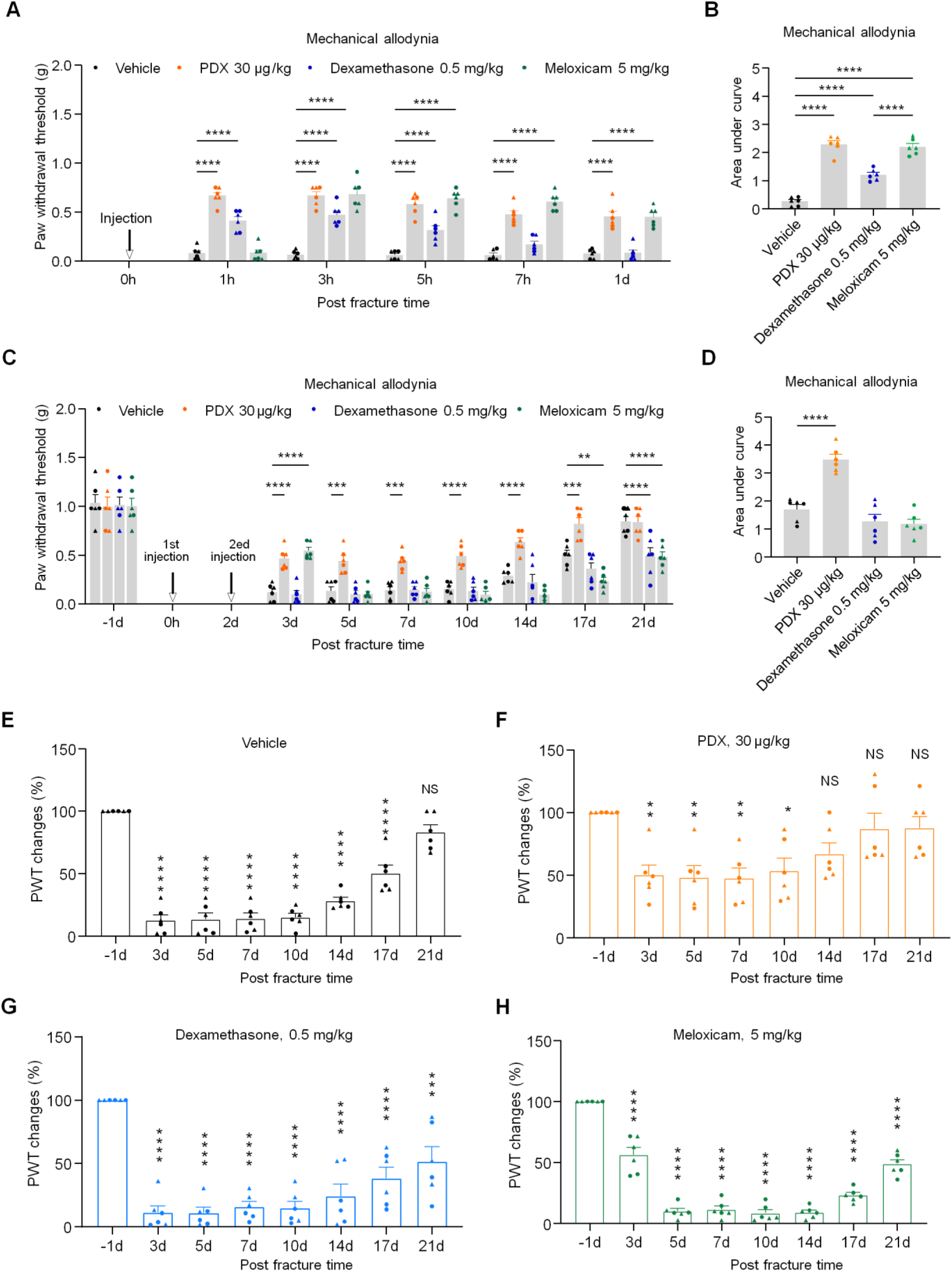
Effects of PDX, dexamethasone, and meloxicam on acute fPOP and the resolution of fPOP in CD1 mice. **(A, B**) Acute effects after the first i.v. injection of PDX (30 μg/kg), dexamethasone (0.5 mg/kg), meloxicam (5 mg/kg), or vehicle (PBS, 100 μl) given immediately before the fracture surgery. The data are shown as PWT (A) and AUC of PWT (B). (**C-D**) Sustained effects of the drugs on the resolution of fPOP following the 2^nd^ injection given 48h after tibial surgery. The data are shown as PWT (C) and AUC of PWT (D). (**E-H**) Time course of fPOP showing distinct recovery of fPOP following different pain treatments. The data are shown as percentage of pre-injury baseline PWT after the 2^nd^ injection of vehicle (E), PDX (F), dexamethasone (G), and meloxicam (H). All the data are represented as mean ± SEM and statistically analyzed by two-way ANOVA with Bonferroni’s post hoc test (A, C) and one-way ANOVA with Bonferroni’s post hoc test (B, D, E-H). \**P* < 0. 05, \*\**P* < 0. 01, \*\*\**P* < 0.001, \*\*\*\**P* < 0.0001; *n* = 6 mice of equal sex; ▴ male, ● female.

Recovery profiles further showed full resolution by day 21 in vehicle controls (Figure 2E), accelerated recovery by day 14 with PDX (Figure 2F), and delayed or absent recovery by day 21 with dexamethasone or meloxicam (Figure 2G,H). Thus, unlike current anti-inflammatory treatments that can delay pain resolution, PDX both relieves and accelerates recovery from fPOP.

### PDX is endogenously produced in muscle tissue at the fracture site

We performed lipidomic analysis to compare the endogenous production of PD1 and PDX in muscle tissue surrounding the tibial bone (Figure 3A-F). Lipid mediator quantification was carried out using liquid chromatography and tandem mass spectrometry (LC-MS/MS) on SCIEX Triple Quad 7500. Lipid mediators were identified by matching retention time and prominent ions in their MS-MS to those of authentic Serhan Lab Standards for PDX (Supplemental Figure 2) and PD1 (Supplemental Figure 3). Our results showed the presence of PDX (Figure 3A-B) and PD1 (Figure 3D-E) in the muscle tissue. Notably, the endogenous PDX levels were much higher than those of PD1/NPD1 both in naïve animals without surgery (Figure 3C, *P* = 0.106) and in animals with bone fracture (*P* < 0.05, Figure 3C and 3F, Supplemental Table 1). Compared to naïve animals (<50 pg/sample), PDX was further elevated after bone fracture (near 100 pg/sample, Figure 3C and 3F). Strikingly, PDX levels were 10 times greater than PD1 in muscle samples (Supplemental Table 1). We also identified greater levels of PDX compared to PD1 in the spleen, an immune organ (Supplemental Table 2). The ratio of PDX in total free fatty acids was also greater in muscle than spleen tissues (Figure 3G). Furthermore, PDX levels were among the highest of all the detected SPMs in muscle and spleen tissues (Supplemental Tables 1–2), underscoring its unique role in inflammation resolution and pain modulation.

**Figure 3.**
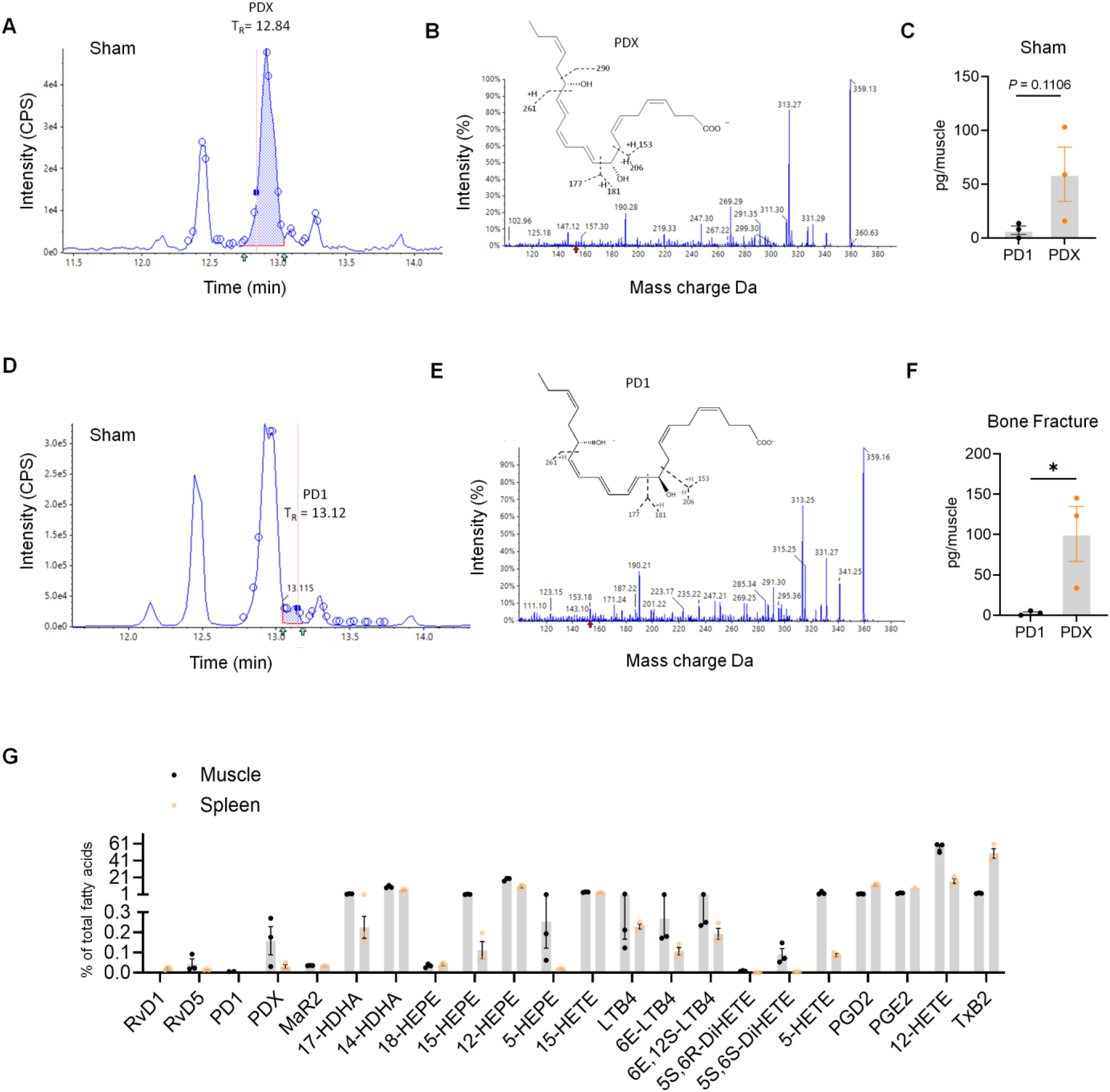
Lipidomic analysis reveals distinct production of PDX and PD1 at the fracture site of CD1 mice. (A, D) Chromatogram showing retention times for PDX (A) and PD1 (D) in bone fracture or sham muscle tissues. (B, E) Mass spectrum showing fragmentation pattern, as well as prominent ions and chemical structure of PDX (B) and PD1 (E). (C, F) Quantification of PDX and PD1 levels in sham muscle tissue (C; n = 3) or bone fracture muscle tissue (F; n = 3). (G) Normalized lipid mediator levels as % of total fatty acid level in muscle and spleen (n = 3) tissues of mice with bone fracture. All the data are represented as mean ± SEM and statistically analyzed by unpaired t-test (C, F). \**P* < 0.05. CD1 male mice were given either sham surgery or tibial bone fracture surgery, and muscles and spleen tissues were collected 3 days after surgery and stored in PBS. Lipid mediator quantitation was carried out using liquid chromatography and tandem mass spectrometry (LC-MS/MS) on SCIEX Triple Quad 7500. Lipid mediators were identified by matching retention and prominent ions in their MS-MS to those of authentic Results expressed as pg/60 mg spleen tissue. § = not identified.

### PDX binds and activates GPR37 to induce Ca²⁺ signaling

Because GPR37 serves as a receptor for NPD1 (18,19) and PDX is structurally related, we examined whether PDX also targets GPR37. Homology modeling and molecular dynamics simulations predicted stable PDX– GPR37 interactions involving ARG442, ASN531, and ARG401, with consistent binding stability over 100 ns (RMSD 4–6 Å; Figure 4A,B). To test GPR37-mediated signaling, we measured intracellular Ca²⁺ in HEK293T cells expressing human GPR37. PDX (30 nM) evoked robust Ca²⁺ increases only in GPR37⁺ cells (EC₅₀ ≈ 23.5 nM), reversible on washout, and ATP produced expected responses (Figure 4C–F). Lipid overlay assays further confirmed direct binding of PDX and NPD1 to GPR37 (Figure 4G–I). Native HEK cells showed minimal responses even at 30 nM, consistent with low endogenous GPR37 expression (Supplemental Figure 8A–C). Given high Gpr37 expression in peritoneal macrophages (19), we evaluated Ca²⁺ responses in WT and *Gpr37*⁻^/^⁻ macrophages. PDX triggered rapid, dose-dependent Ca²⁺ elevations in WT macrophages (EC_50_ ≈ 4.2 nM)—notably more sensitive than HEK293T cells—and failed to induce responses in *Gpr37*⁻^/^⁻ cells (Figure 4J,K). Together, these data identify GPR37 as a functional receptor for PDX and demonstrate GPR37-dependent Ca²⁺ signaling in both engineered and native immune cells.

**Figure 4.**
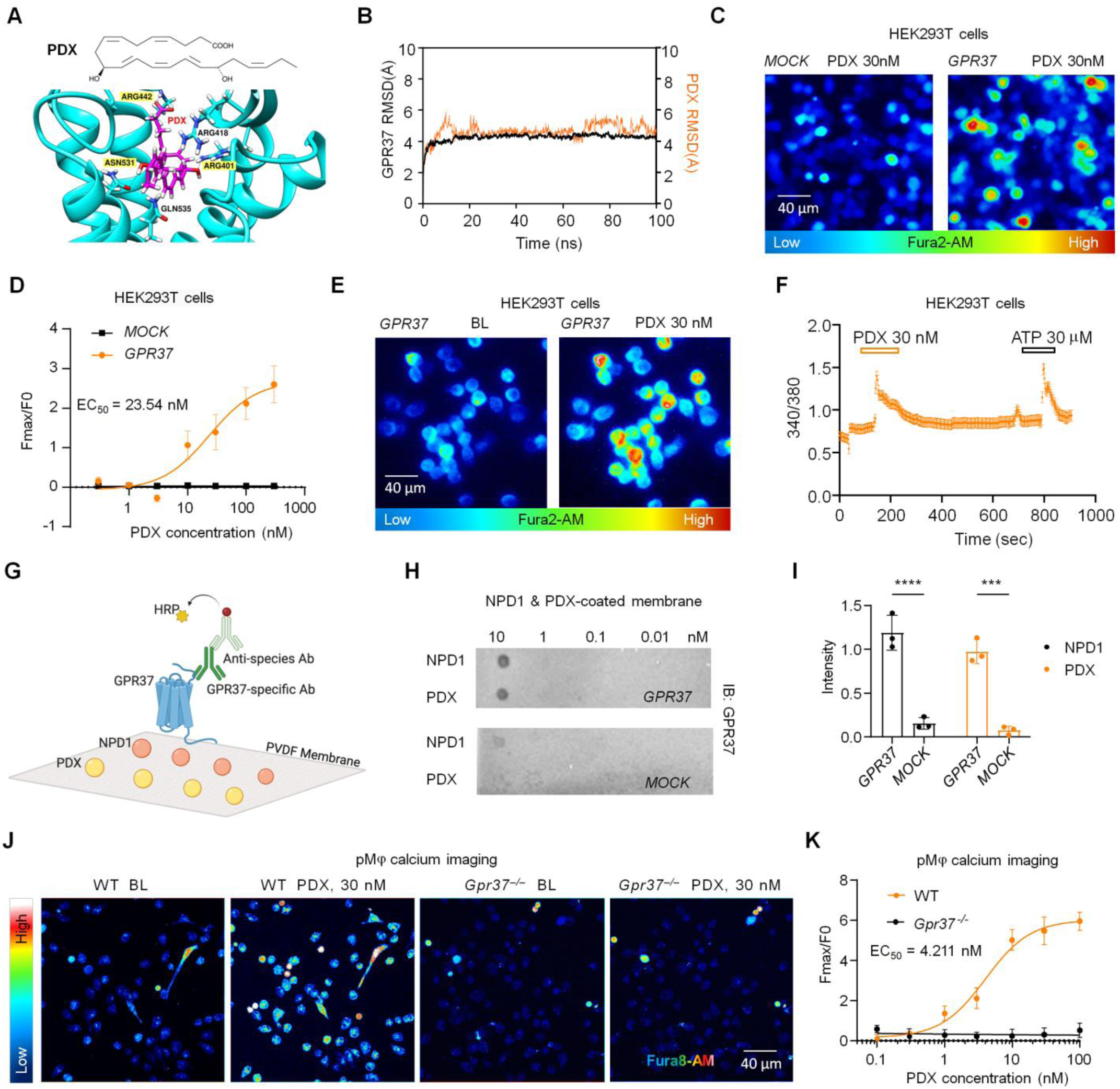
PDX directly binds GPR37 and increases calcium influx in HEK293T cells and peritoneal macrophages (pM𝜑s) from C57BL/6 mice. **(A**) Structural model of human GPR37 (blue) in complex with PDX (magenta). (**B**) Molecular dynamics simulation showing root mean square deviation (RMSD) of the GPR37–PDX complex (magenta) versus GPR37 alone (blue) over 100 ns, indicating stable ligand–receptor interaction. (**C**) PDX (30 nM) evoked calcium influx in GPR37-transfected HEK293T cells but not in mock-transfected controls. (**D**) Dose-response curve of PDX-induced calcium signaling in GPR37-expressing HEK293T cells versus mock controls (EC_50_ = 23.54 nM; n = 16 reads from 4 cultures). (**E** and **F**) Representative traces (E) and quantification (F) demonstrate enhanced calcium responses to PDX (30 nM) and ATP (30 µM) in GPR37-expressing HEK293T cells (n = 27 cells, 3 cultures). (**G**) Dot-blot schematic for assessing binding of GPR37 to PDX and NPD1 using PVDF membranes coated with ligands at graded concentrations. (**H**) Representative blot of PDX and NPD1-coated PVDF membranes incubated with lysates from HEK293T cells with and without GPR37 expression. (**I**) Quantification of dot intensity in HEK293T cell lysates with and without GPR37. n = 3 repeats. (**J** and **K**) Calcium imaging in pM𝜑s showing traces (J) and quantification (K) of calcium responses following PDX (30 nM) treatment in pM𝜑 cultures prepared from WT or *Gpr37*^−/−^. Note that PDX induces dose-dependent responses in WT pM𝜑s but has no effects in *Gpr37*^−/−^ pM𝜑s. EC_50_ of the PDX-induced calcium response is 4.21 nM. n = 9 cultures from 3 mice. The Data are expressed as means ± SEM. Statistics: two-way ANOVA with Tukey’s post hoc test (I). \*\*\**P* < 0.001, \*\*\*\**P* < 0.0001. Scale bars: 40 μm (C, E, J).

### GPR37 activation accelerates fPOP resolution and mediates PDX-induced pain relief

To define the role of GPR37 in postoperative pain, we compared WT and *Gpr37*^−/−^ mice (C57BL/6). In WT mice, mechanical allodynia resolved by ∼5 weeks and cold hypersensitivity by ∼6 weeks (Figure 5A,B), consistent with longer fPOP duration in C57BL/6 than CD1 mice. In contrast, *Gpr37*^−/−^ mice showed markedly prolonged pain, with full recovery only after ∼5 months and significantly lower PWT from 5 weeks to 4 months (*P* < 0.0001). Thus, GPR37 is essential for fPOP resolution. We also observed sex differences: female *Gpr37*^−/−^ mice exhibited greater cold pain than males from day 3 to month 3 (*P* < 0.01; Figure 5C). To test whether GPR37 mediates PDX action, we administered PDX (100 ng, i.v.) at 3 weeks after fracture. PDX reduced mechanical, thermal, and cold hypersensitivity in WT mice, but had no effects in *Gpr37*^−/−^ mice (*P* < 0.01 vs. WT; Figure 5D–F). These results show that GPR37 promotes resolution of postoperative pain and is required for PDX-induced analgesia.

**Figure 5.**
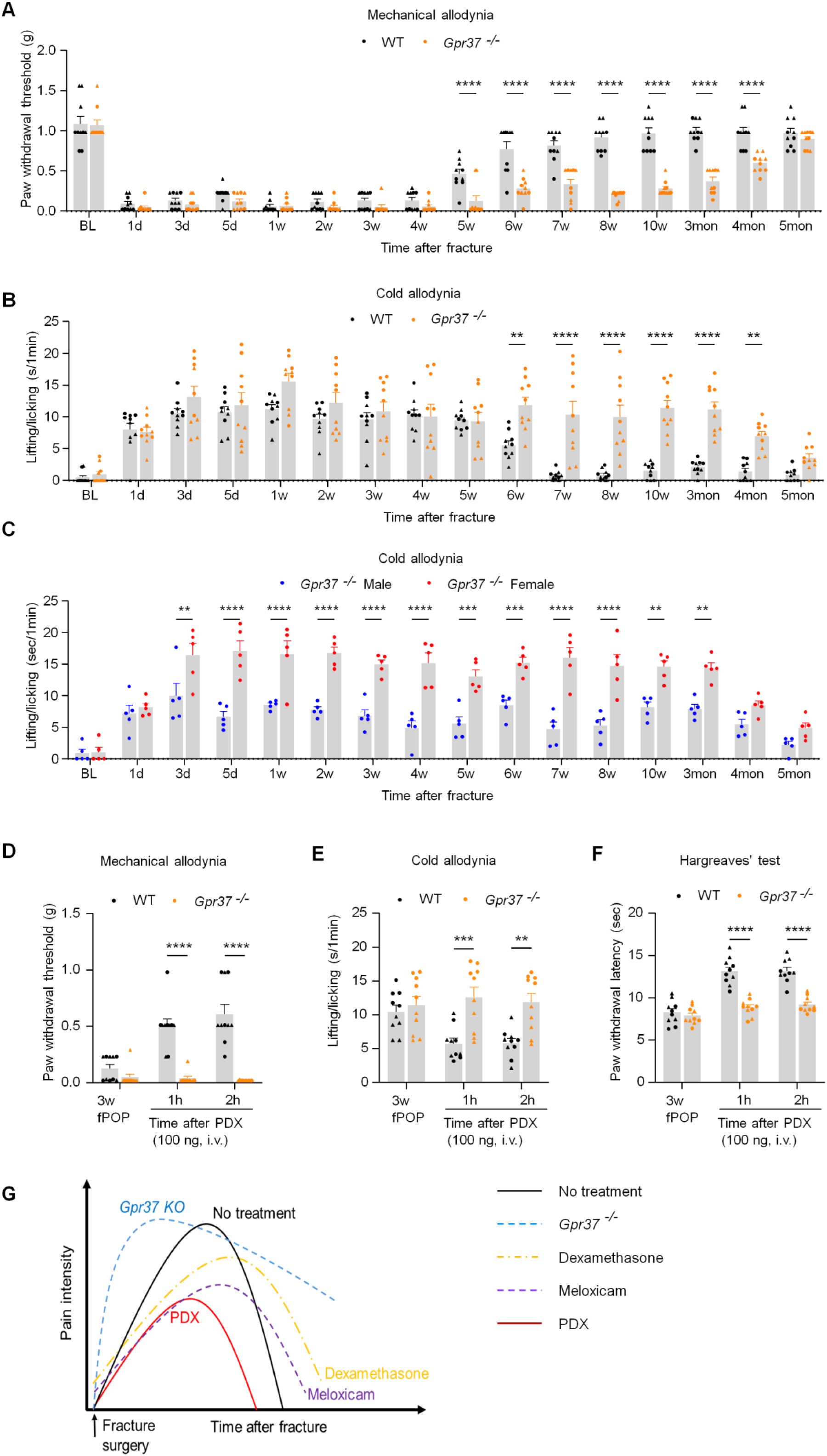
GPR37 mediates PDX-induced pain relief and contributes to the resolution of fPOP in C57BL/6 mice. (**A, B**) Time course of mechanical allodynia (von Frey test, A) and cold allodynia (acetone test, B) following tibial fracture in WT and *Gpr37*^−/−^ mice. *Gpr37*^−/−^ mice showed delayed recovery, indicating impaired resolution of postoperative pain. (**C**) Sex-specific analysis of cold allodynia in *Gpr37*^−/−^ mice reveals persistent cold pain in females up to 3 months post-fracture, whereas males exhibit partial recovery. (**D-F**) PDX treatment (100 ng, i.v.) reduced fPOP only in WT mice but not in *Gpr37*^−/−^ mice, as assessed in von Frey test (**D**), Acetone test (**E**), and Hargreaves’ test (**F**) in WT and *Gpr37*^−/−^ mice. Data are represented as mean ± SEM and statistically analyzed by two-way ANOVA with Bonferroni’s post hoc test (**A-F**). \*\**P* < 0. 01, \*\*\**P* < 0.001, \*\*\*\**P* < 0.0001; *n* = 10 (A-B, D-F), *n* = 5 (C); ▴ male, ● female. **(G)** Schematic illustration of pain resolution following fracture surgery in WT mice under different treatment conditions: 1) no treatment, 2) NSAID (meloxicam), 3) steroid (dexamethasone), 4) PDX treatment, and 5) *Gpr37*^−/−^ mice. Notably, PDX accelerates pain resolution, whereas NSAID and steroid treatments delay resolution. Also note that *Gpr37*^−/−^ mice fail to resolve pain.

### Pathway analysis reveals macrophage/neutrophil signaling and wound healing upregulated by PDX treatment in fPOP

Primary sensory neurons, glial cells, and immune cells in the DRG play an important role in the pathogenesis of pain (32, 33). We collected L_4_-L_5_ DRG tissues from mice 3 days after fracture with and without PDX treatment and conducted bulk RNA-sequencing (RNAseq). The canonical pathway analysis showed significant gene changes related to macrophage and neutrophil function and cytokine responses (Figure 6A). PDX upregulated many genes related to macrophage function, including pathogen recognition (*Camp, Cybb*), phagocytosis (*Itgam*), inflammation regulation (*Cxcr2, Ccr1*), wound healing (*Ppbp*), and immune response, especially S100 family genes (*S100a9, S100a8*) and pro-resolution gene (*Frp2*) encoding the SPM lipoxin A4 receptor ALX/FPR2 (34, 35) (Figures 6B-H). These results suggest that PDX might trigger an immune activation in the DRG of mice with fPOP, involving macrophages and neutrophils.

**Figure 6.**
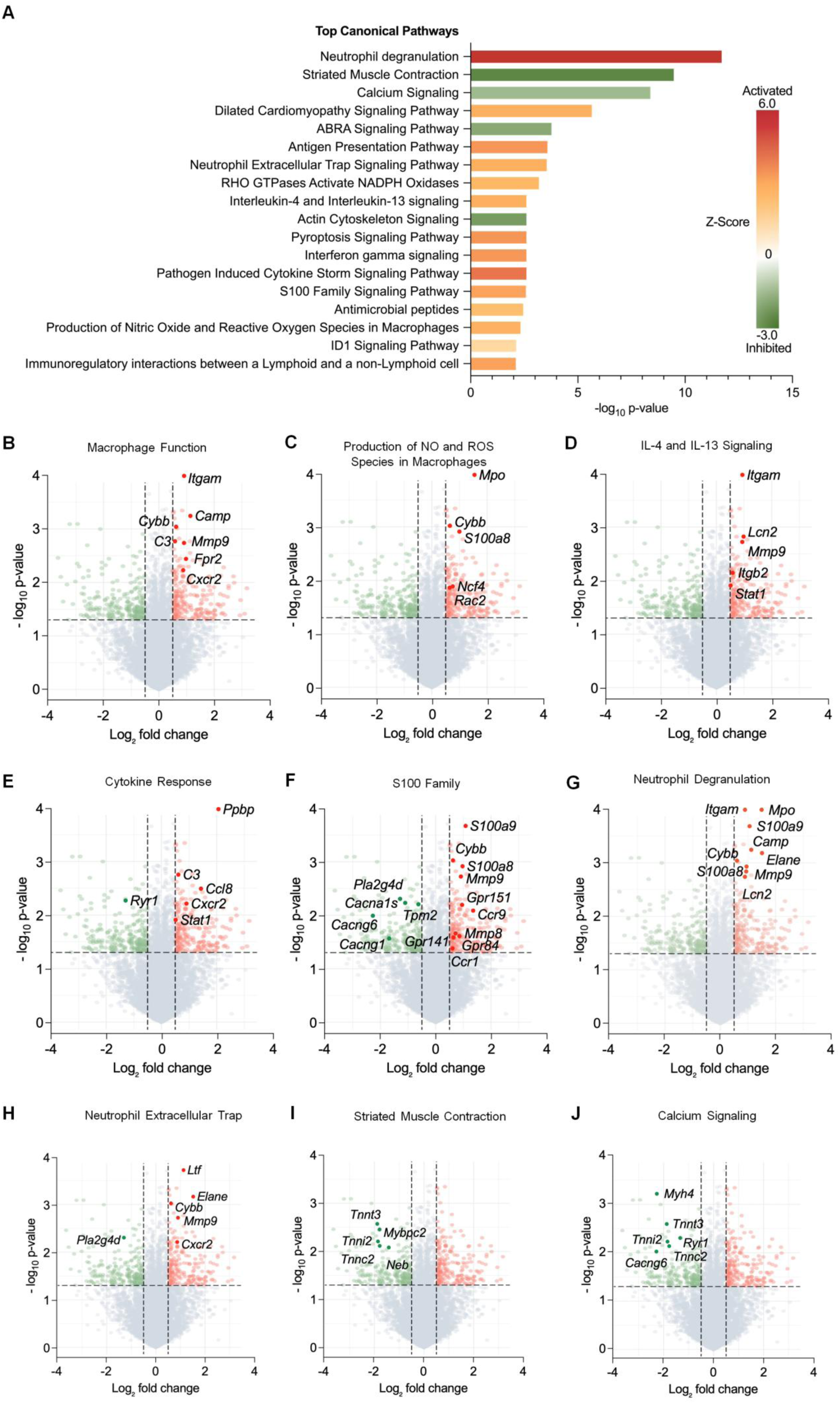
Bulk RNA-sequencing shows the PDX effects on multiple pathways in DRG of mice with fPOP in CD1 mice. (**A**) Bulk RNA-sequencing data analysis showing the top canonical pathways in fracture + PDX versus fracture + vehicle group. All the pathways show statistical significance of *P* < 0.01 with both upregulations and downregulations as shown in Z-Score. (**B-J)** Volcano plots for differentially expressed genes. Red dots denote the up-regulated genes, and green dots denote the down-regulated genes with > 50% change and *P* < 0.05. The activated pathways are related to macrophage function (B), production of NO and ROS species in macrophage genes (C), IL-4 and IL-13 signaling genes (D), pathogen-induced cytokine genes (E), S100 family genes (F), neutrophil degranulation genes (G), and neutrophil extracellular trap genes (H). The inactivated pathways are related to striated muscle contraction genes (I) and calcium signaling genes (J). *n* = 6 mice (3 males and 3 females) per group.

### PDX regulates pro- and anti-inflammatory cytokine expression in macrophages via GPR37

To understand the role of PDX in regulating the expression of key inflammatory cytokines in pM𝜑s cultures we employed quantitative PCR analysis. We cultured pM𝜑s from WT and *Gpr37*^−/−^ mice, treated the cells with PDX (30 nM) and/or LPS (1 μg/ml) for 12 hours, and measured the mRNA levels of pro-inflammatory cytokines (*Il1b* and *Tnf*) and anti-inflammatory cytokine *Il10* (Supplemental Figure 9A). In cultured WT pM𝜑, PDX treatment increased *Il10* mRNA levels and inhibited the LPS-induced increase in *Il1b* and *Tnf* mRNA expression (Supplemental Figure 9B). However, PDX treatment did not reverse the LPS-induced changes in *Il1b*, *Tnf*, and *Il10* in *Gpr37*^−/−^ pM𝜑s (Supplemental Figure 9C). Additionally, ELISA was used to detect the secretion of IL-1β, TNF-α, and IL-10 in the pM𝜑s from WT and *Gpr37*^−/−^ mice (Supplemental Figure 9D). PDX reversed the LPS-induced secretion of pro-inflammatory cytokines (IL-1β, TNF-α) by pM𝜑s. PDX treatment alone also increased the secretion of IL-10 (Supplemental Figure 9E-G). These effects of PDX were lost in *Gpr37*^−/−^ pM𝜑s (Supplemental Figure 9H-J). Collectively, our findings suggest a important role of PDX in regulating pro- and anti-inflammatory cytokine expression in macrophages through GPR37.

### PDX accelerates macrophage phagocytosis via GPR37 and Ca^2+^ signaling

Because NPD1 promotes macrophage phagocytosis (18), we tested whether PDX does the same. pH-sensitive zymosan assays showed that PDX (30 nM, 1 h) significantly accelerated and increased phagocytosis in mouse peritoneal macrophages, with higher phagocytic rates at 15–45 min and near-complete uptake by 60 min (Figure 7A–C). PDX similarly enhanced phagocytosis in THP1 human macrophages (Figure 7D–F).

**Figure 7.**
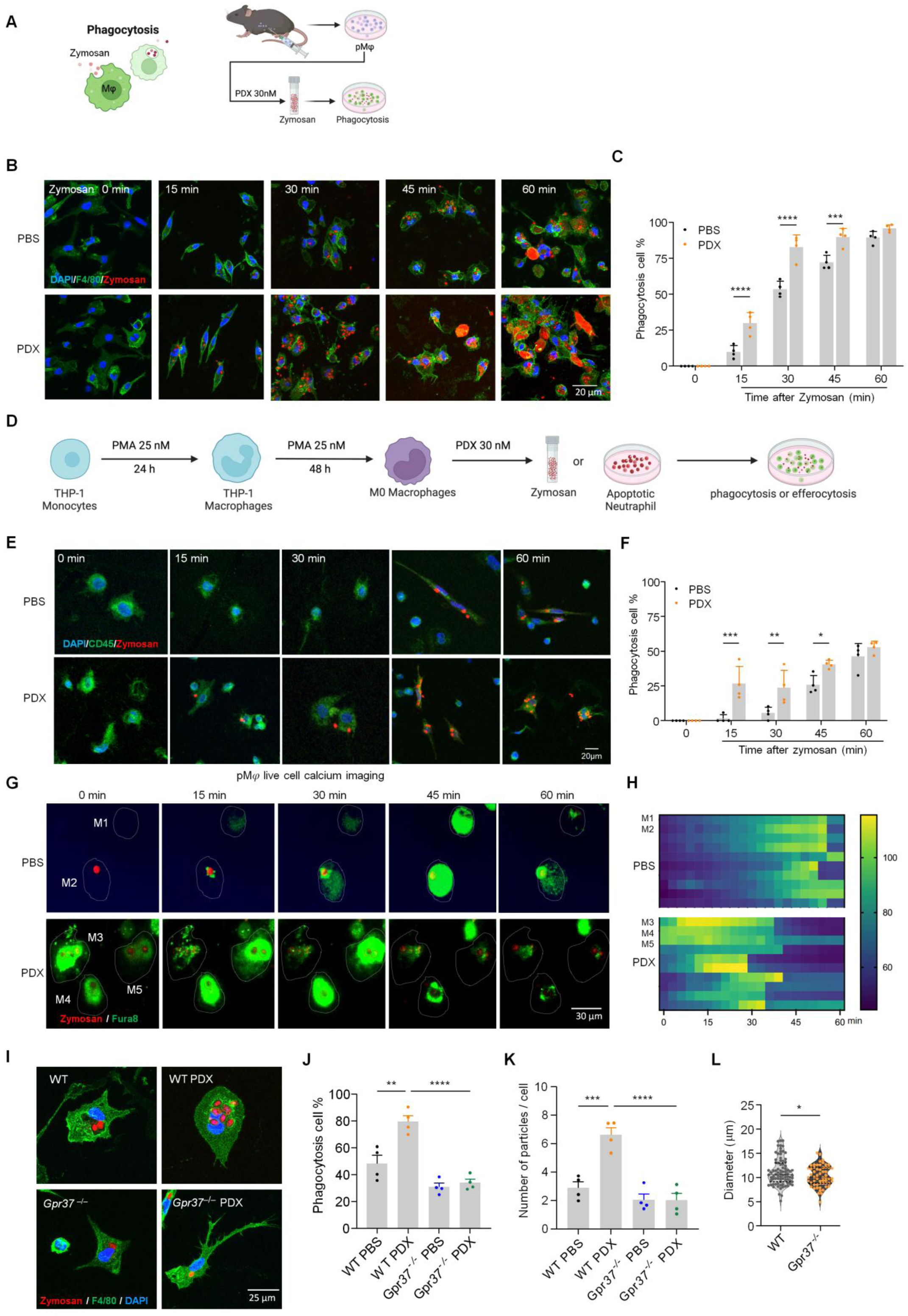
PDX increases macrophage phagocytosis in pM𝜑s of C57BL/6 mice and THP1-derived macrophages via calcium signaling. **(A**) Schematic of zymosan phagocytosis assay in primary pMΦs. pHrodo Red zymosan particles (red; activated in acidic phagosomes) were added 1 h before imaging. (**B**) Representative fluorescent images showing time-dependent zymosan uptake in WT pMΦs labeled with F4/80 (green) at 0, 15, 30, 45, and 60 min after PBS or PDX treatment. (**C**) Quantification of phagocytic pMΦs (% cells containing >1 particle). (**D**) Schematic of zymosan phagocytosis in THP1 culture. THP1 monocytes were differentiated into macrophages with PMA (25 nM, 48 h). (**E** and **F**) Time-lapse images (E) and quantification (F) of zymosan uptake in THP1 macrophages (CD45+, green) over 60 min. PDX accelerated phagocytosis relative to PBS. **(G)** Calcium imaging of pMΦs during phagocytosis, showing increased calcium influx with PDX versus PBS at multiple timepoints. Fura-8 AM (green) was used for Ca²⁺ detection; zymosan shown in red. pM𝜑s were indicated by the white dotted circle outlines and labeled as M1-M5. **(H)** Heat map shows time-dependent calcium influx of pM𝜑s after PBS (n = 10 cells) or PDX (n = 10 cells) incubation, including M1-M5 presented in panel G. This experiment was conducted in 3 cultures. (**I-K**) PDX-induced zymosan phagocytosis of pM𝜑s (F4/80^+^, green) from WT and *Gpr37*^−/−^ mice, as shown by images at 0 and 30 min (I) and quantification of percentage of positive cells with phagocytosis (J) and zymosan particle number in each cell (K). *n* = 4 cultures. (L) Cell diameters of WT and *Gpr37*^−/−^ pM𝜑s. *n* = 83 cells from 4 cultures from WT mice; *n* = 58 cells from 4 cultures from KO mice. Data are mean ± SEM and statistically analyzed by 2-way ANOVA with Bonferroni’s post hoc test. \**P* < 0.05, \*\**P* < 0.01, \*\*\**P* < 0.001, \*\*\*\**P* < 0.0001; *n* = 4 cultures/condition (12 mice total; mixed sex, 7C and 7D). *n* = 4 cultures/condition (THP1 cells in 7E and 7F).

Live Ca²⁺ imaging revealed that PDX advanced intracellular Ca²⁺ elevations coincident with zymosan uptake, whereas vehicle produced delayed Ca²⁺ signals and slower phagocytosis (Figure 7G,H; Videos 1–2), indicating Ca²⁺-dependent phagocytosis. PDX failed to enhance phagocytosis in *Gpr37*^−/−^ macrophages, which also showed reduced baseline uptake and smaller cell size (Figure 7I–L), demonstrating that GPR37 is required for PDX-driven phagocytosis and macrophage growth.

Mechanistically, PDX-induced phagocytosis was abolished by Ca²⁺ chelation (BAPTA-AM) and significantly inhibited by pertussis toxin, a Gβ/γ blocker, and a PI3K/AKT inhibitor (Supplemental Figure 10B), implicating GPR37–Gi/o–Ca²⁺–PI3K/AKT signaling. Thus, PDX promotes macrophage phagocytosis through GPR37-dependent Ca²⁺ signaling and downstream Gi/βγ/PI3K pathways.

### PDX enhances macrophage efferocytosis via GPR37 and Ca^2+^ signaling

Macrophage efferocytosis of apoptotic cells is essential for tissue repair and inflammation resolution (36, 37). To test whether PDX promotes this process, we exposed pMφs to apoptotic, fluorescently labeled neutrophils. PDX significantly increased efferocytosis at 2 and 3 hours compared to vehicle (Figure 8A–C). PDX similarly enhanced efferocytosis in THP1 macrophages (Figure 8D–F), consistent with its role in promoting resolution. PDX-induced efferocytosis was blocked by Ca²⁺ chelation (BAPTA-AM) and inhibited by pertussis toxin, a Gβ/γ blocker, and PI3K/AKT inhibition, indicating dependence on GPR37–Gi/o–Ca²⁺–PI3K signaling (Supplemental Figure 10C). To assess clearance in vivo, we quantified apoptotic neutrophils at fracture sites. PDX enhanced efferocytosis in WT mice but not in *Gpr37*^−/−^ mice (Figure 8G–I). PDX also modestly improved macrophage survival in WT mice, *Gpr37*^−/−^ mice showed reduced macrophage survival (Figure 8J,K). Thus, PDX promotes efferocytosis and macrophage survival through GPR37-dependent Ca²⁺ and Gi/βγ/PI3K-AKT signaling *in vitro* and *in vivo*.

**Figure 8.**
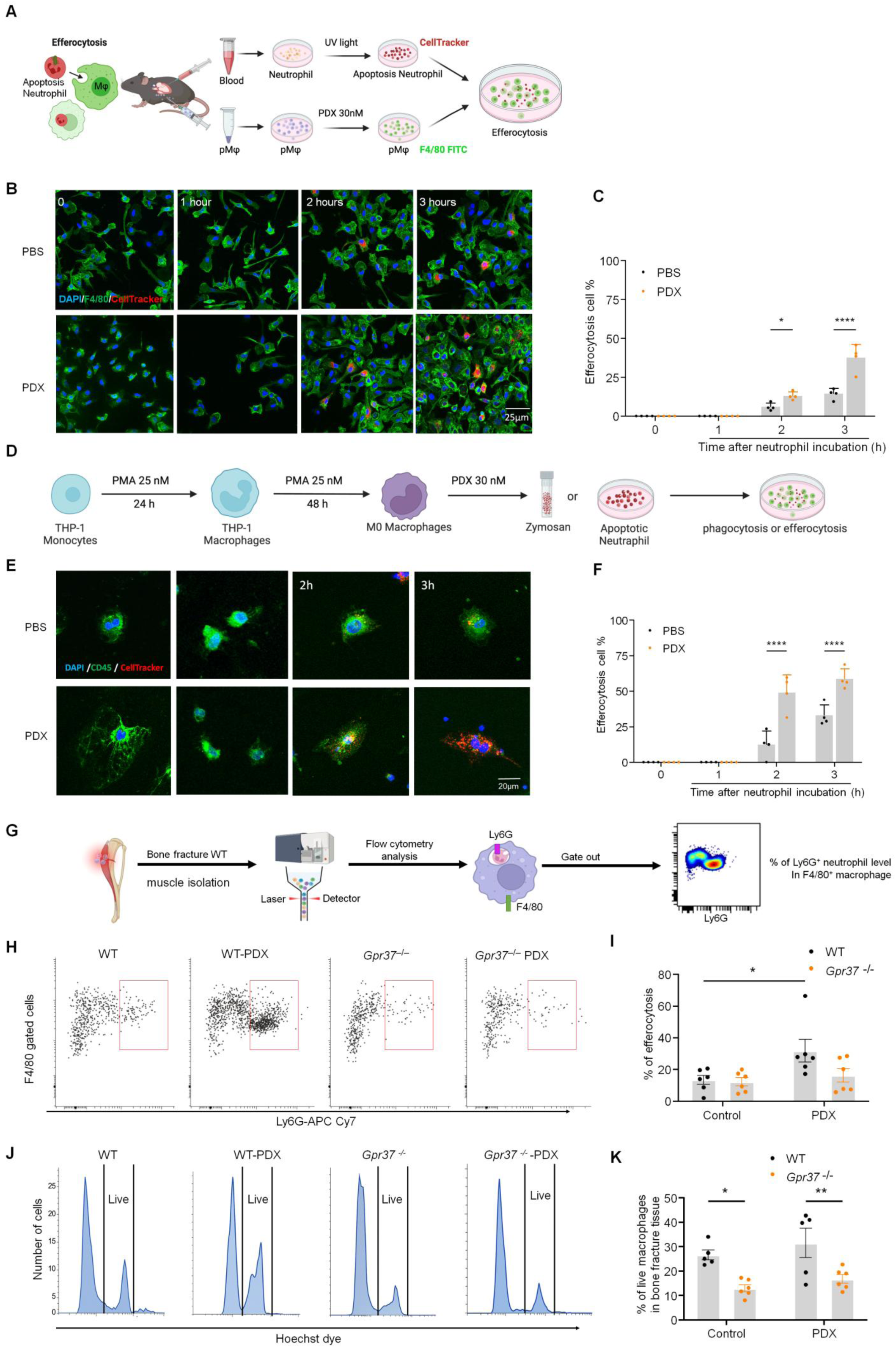
PDX accelerates macrophage efferocytosis of neutrophils in C57BL/6 mice via GPR37. (**A**) Schematic of neutrophil efferocytosis assay in pM𝜑 culture. Neutrophils from mouse whole blood were exposed to UV light for 10 min to induce apoptosis and then labeled with CellTracker (Red). The CellTracker labeled apoptotic neutrophils were incubated with pM𝜑, which were labeled with F4/80 (Green). (**B**) Fluorescent images showing time-dependent efferocytosis in WT pM𝜑s at 0, 1, 2, and 3 hours (h) after PBS and PDX treatment. Scale bar: 25 μm. (**C**) Quantification of percentage of pM𝜑s with efferocytosis. The green macrophages containing neutrophils or cell debris were regarded as positive cells. (**D**) Schematic of neutrophil efferocytosis in THP1 culture. PMA (25 nM, 48 h) was used to stimulate cell differentiation from monocytes to M𝜑s. (**E** and **F**) Images (E) and quantification (F) of THP1 M𝜑 efferocytosis of apoptotic neutrophils (Red, CellTracker) at 0, 1, 2, and 3 h. Scale bar, 20 μm. (**G**) Schematic of neutrophil efferocytosis assay in bone fracture muscle tissue by pM𝜑. (**H, I**) Effects of PDX treatment on neutrophil efferocytosis activity (H) and macrophage abundance (I) in the bone fracture tissue collected from WT and *Gpr37*^−/−^ mice (100 ng, I.V., n = 6). (**J, K**) Effects of PDX treatment on F4/80-positive macrophage survival ratio using flow cytometry analysis from WT and *Gpr37*^−/−^ mice (100 ng, i.v., n =6). Data are mean ± SEM. Statistics: one-way ANOVA with Bonferroni’s post hoc test (B and F) and two-way ANOVA with Bonferroni’s post hoc test (I and K). \**P* < 0.05, \*\**P* < 0.01, \*\*\**P* < 0.001, \*\*\*\**P* < 0.0001.

### PDX inhibits C-fiber reflex and DRG neuron Ca^2+^ responses under fPOP

RNA-seq of DRG from fracture mice showed that PDX down-regulated muscle contraction and calcium-signaling pathways (Figure 6A, I, J), suggesting modulation of muscle nociception—an important component of postoperative pain. We assessed nociceptive C-fiber reflexes via biceps femoris electromyogram (EMG, Figure 9A,B). Tibial fracture lowered reflex threshold and latency and increased amplitude and firing frequency (*P* = 0.0376; *P* < 0.001), consistent with sensitization (Figure 9C–F). Peri-sciatic PDX (20 ng, 30 min) reversed these changes, and intraperitoneal PDX (150 ng) similarly reduced reflex amplitude and frequency (Supplemental Figure 11A–D). Thus, PDX blocks fracture-induced C-fiber reflex facilitation.

**Figure 9.**
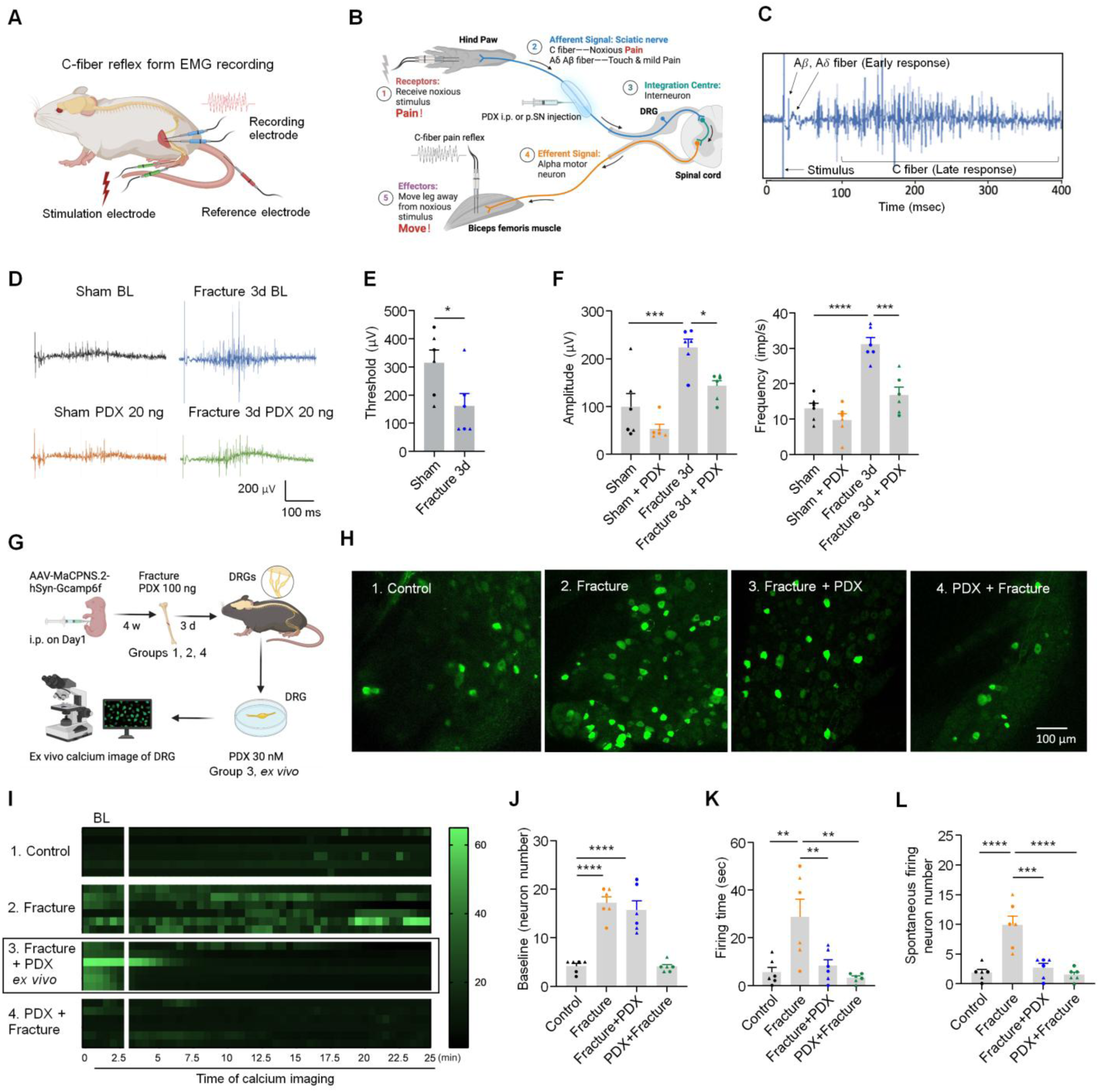
PDX inhibits the C-fiber reflex in vivo and DRG spontaneous neuronal activity ex vivo after fracture in CD1 mice. (**A,B**) Schematic illustrating EMG recording of the C-fiber reflex in the biceps femoris, including electrode placement (A) and the afferent–efferent spinal reflex arc (B). (**C**) Representative EMG traces showing A-fiber mediated early responses and C-fiber mediated late responses. (**D**) C-fiber reflex recordings in four groups: sham or fracture surgery with PBS or PDX. PDX (20 ng, 20 μl) was administered by peri-sciatic nerve (p.SN.) injection. EMG was recorded before (baseline) and 30 min after treatment. (**E**) Thresholds 3 days after sham and fracture surgery. (**F**) EMG amplitude (left) and frequency (right) from 4 groups in D. EMG data were quantified on post-surgical day 3. (**G**) Schematic of ex vivo whole-mount DRG calcium imaging in four treatment groups: sham (Group 1), fracture (Group 2), fracture with ex vivo PDX bath application (30 nM; Group 3), and fracture with in vivo PDX pretreatment (100 ng i.v.; Group 4). AAV-MaCPNS.2-hSyn-Gcamp6f was delivered at P1, and ipsilateral L3–L5 DRG were collected on day 3 after fracture for imaging using confocal microscopy. (**H-I**) Representative Ca²⁺ traces (H) and heatmaps (I) demonstrate that fracture induces spontaneous DRG neuronal activity, which is prevented by in vivo PDX and reversed by ex vivo PDX (n=6 neuron/group). (**J-L**) Quantification of calcium signal in DRG neurons, showing number of neurons per DRG with spontaneous activity at the baseline (J), firing time of DRG neurons for all the firing events in each DRG (K), and the number of spontaneously discharging neurons in each DRG (L). Data are mean ± SEM. Statistics: one-way ANOVA with Bonferroni’s post hoc test. \**P* < 0.05, \*\**P* < 0.01, \*\*\**P* < 0.001, \*\*\*\**P* < 0.0001; *n* = 6 mice/group; ▴ male, ● female.

To examine sensory neuron activity, we performed *ex vivo* Ca²⁺ imaging of L3/4 DRG expressing GCaMP6f (AAV-MaCPNS.2-hSyn-GCaMP6f; Figure 9G). Fracture increased baseline Ca²⁺ activity by day 3, including longer spontaneous discharge and more active neurons (Figure 9H–L). Both pre- and post-treatment with PDX markedly reduced spontaneous Ca²⁺ events and the number of hyperactive neurons (Figure 9I–L). These results indicate that fracture induces DRG neuron hyperexcitability and enhanced C-fiber reflexes, whereas PDX normalizes neuronal activity and suppresses nociceptive reflex facilitation.

### PDX inhibits TRPA1/TRPV1-induced pain and neurogenic inflammation

Transient receptor potential ion channels A1 and V1 (TRPA1 and TRPV1), expressed by nociceptors, play a crucial role in nociception, and furthermore, SPMs can modulate their activity (16, 38). To test whether PDX reduces TRPA1/V1 channel-evoked pain, we conducted intraplanar injection of TRPA1 agonist AITC (100 μg, i.pl.) or TRPV1 agonist capsaicin (1 μg). PDX markedly suppressed AITC-evoked spontaneous and mechanical pain (*P* < 0.0001; Figure 10A–C), and similarly reduced capsaicin-evoked pain behaviors (Supplemental Figure 12A–C). PDX also diminished AITC- and capsaicin-induced paw edema, as indicated by Evans blue extravasation (*P* < 0.0001; Figure 10D–F; Supplemental Figure 12D–F).

**Figure 10.**
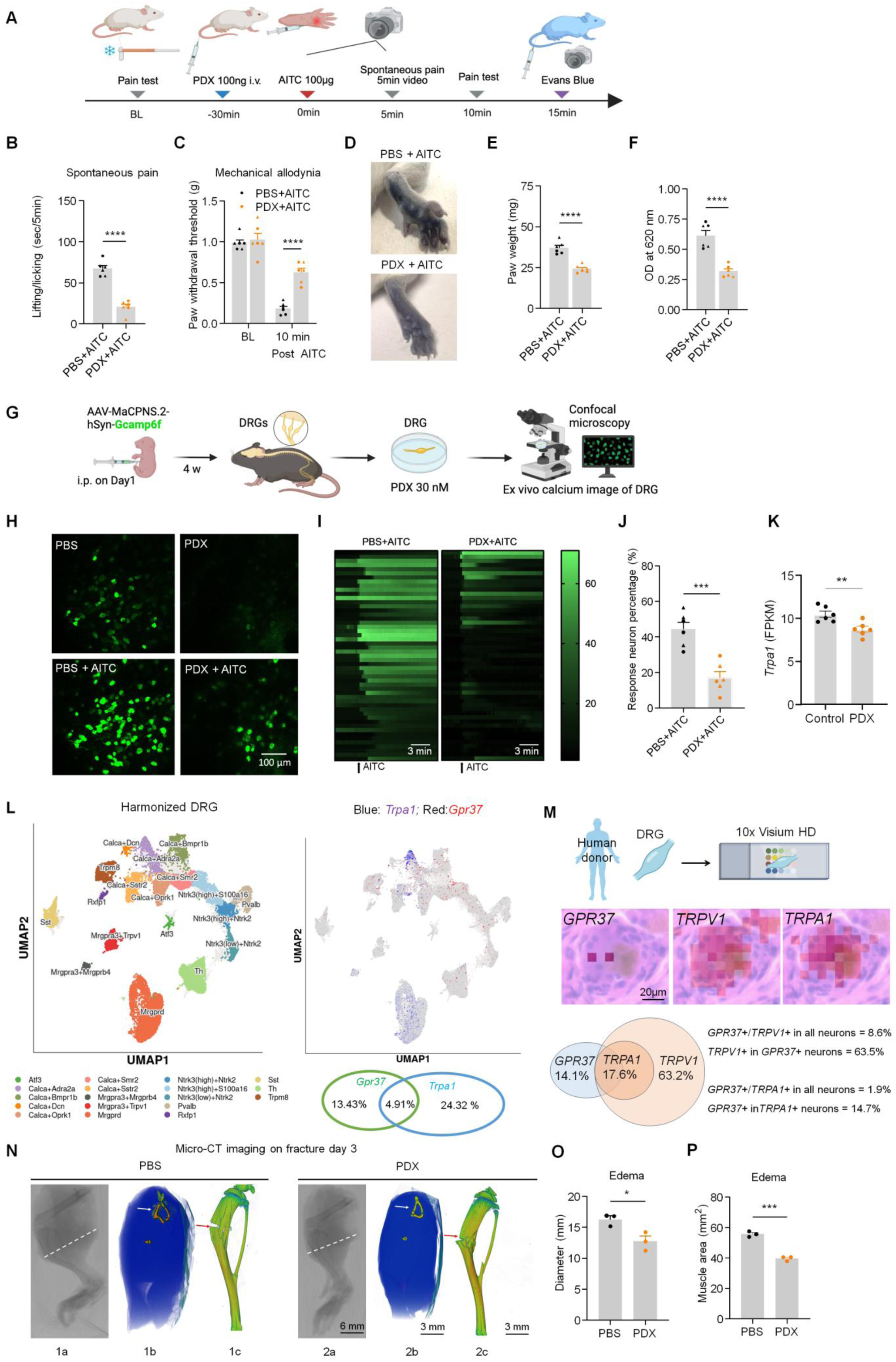
PDX reduces TRPA1-mediated spontaneous pain and neurogenic inflammation in CD1 mice. (**A**) Experimental scheme: mice received PDX 30 min before intraplantar AITC (100 μg); spontaneous pain was recorded, followed by i.v. Evans Blue to assess neurogenic inflammation. (**B**) PDX reduced AITC-evoked spontaneous pain (5 min). (**C**) PDX decreased AITC-induced mechanical allodynia (von Frey, 10 min). (**D**) Representative hind paw images showing AITC-induced edema. (**E, F**) Quantification of edema by paw weight (E) and Evans Blue extravasation (F) with and without PDX (100 ng, i.v.). (**G**) Schematic of *ex vivo* Ca²⁺ imaging in AAV-MaCPNS.2-hSyn-Gcamp6f-labeled neurons. (**H, I**) Representative Ca²⁺ responses (H) and heatmaps (I) showing AITC-evoked neuronal activation. Each group includes 51 DRG neurons. (**J**) PDX decreased the proportion of AITC-responsive DRG neurons. (**K**) Bulk RNA-seq revealed reduced *Trpa1* expression in DRG after fracture in PDX-treated mice. (**L**) Left, sub-cluster annotation map of mouse DRG. Right, UMAP plot showing *Gpr37* (blue) and *Trpa1* (red) expression. Published single-cell RNA-seq data of mouse DRG (60) shows the percentages of *Gpr37* and *Trpa1* expressing neurons and their colocalization. (**M**) Spatial transcriptomics of human DRG showing co-localization of GPR37 with TRPA1 and TRPV1 in human sensory neurons. TRPA1 expression was confined to TRPV1⁺ neurons (1062 neurons from 2 donors). (**N**) Micro-CT of hind limbs 3 days post-fracture showed that PDX reduced edema and improved bone integrity. Coronal and 3-D reconstructions revealed reduced soft-tissue swelling and enhanced tibial bone density at the fracture site. 1c and 2c show 3-D constructed tibial bones, and the red arrows indicate the fracture sites. (**O, P**) Quantification of diameters (O), indicated by white lines in 1a and 2a, and coronal section muscle area (P), indicated in 1b and 2b. Data are mean ± SEM. Statistics: unpaired t-test (B,E,F,J,K,O,P); one-way ANOVA with Bonferroni (C). \**P* < 0.05, \*\*\**P* < 0.001, \*\*\*\**P* < 0.0001; n = 6 mice/group for B,C,E,F,J,K; n = 3 males for O,P. ▴ male, ● female.

In DRG neurons, PDX decreased the proportion of AITC-responsive cells from 44.3% to 16.97% and capsaicin-responsive cells from 20.40% to 5.95% (Figure 10G–J; Supplemental Figure 12G–I). RNA-seq in fracture DRG showed reduced *Trpa1* expression after PDX (*P* < 0.01; Figure 10K). Single-cell and spatial transcriptomics confirmed co-expression of GPR37 with TRPA1 and TRPV1 in mouse and human DRG neurons, with TRPA1 exclusively in TRPV1⁺ cells (Figure 10L,M; Supplemental Figure 12J,K). PDX did not directly alter TRPA1 currents in TRPA1-transfected HEK cells, suggesting indirect neuronal modulation (Supplemental Figure 13A,B).

Finally, Micro-CT analysis showed that PDX pre-treatment significantly reduced fracture-induced hind-limb edema, as indicated by decreased limb diameter and muscle area (*P* < 0.001; Figure 10N–P), and also lessened bone damage (Figure 10O). These findings suggest that PDX confers additional benefit by suppressing TRPA1/TRPV1-mediated neurogenic inflammation and protecting bone integrity.

## Discussion

Earlier results demonstrated the anti-inflammatory actions of PDX, which promotes macrophage polarization through PPARγ-dependent signaling pathways (39). PDX mitigates Type II diabetes via activation of a myokine-liver glucoregulatory axis (25). In the present study, we demonstrated that PDX signals through GPR37 and enhances macrophage phagocytosis/efferocytosis using binding, *in vitro*, and *in vivo* assays, as well as *Gpr37* knockout animals. We have shown the crucial role of GPR37 in mediating the analgesic actions and the pro-resolution functions of PDX, via macrophage and neuronal mechanisms (Figure 11A-B).

**Figure 11.**
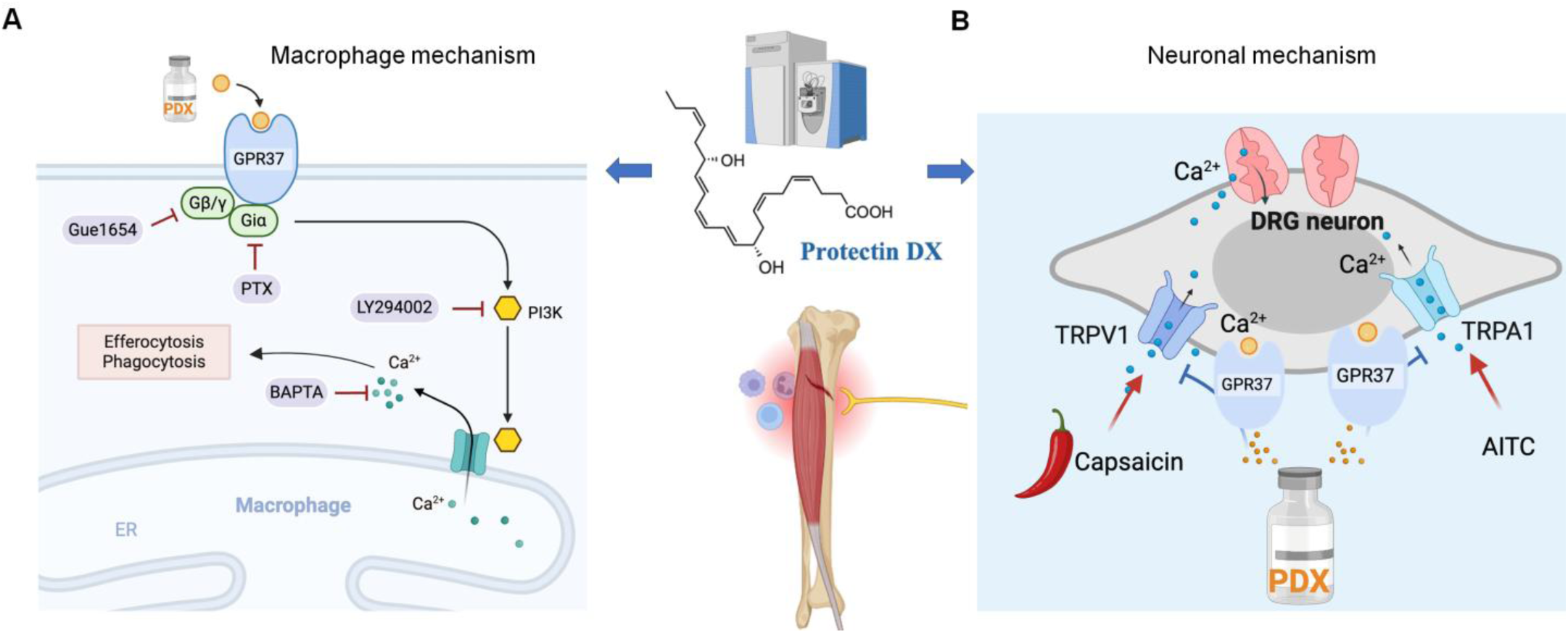
Working hypothesis by which PDX alleviates postoperative pain after tibial bone fracture via macrophage and neuronal mechanisms. **(A)** Schematic illustration of PDX-induced phagocytosis and efferocytosis in pM𝜑s via Giα, Gβ/γ, and PI3K/AKT signaling pathways and intracellular calcium (iCa^2+^) signaling. Macrophage mechanism by which PDX induces phagocytosis/efferocytosis and differentially regulates the expression of pro-inflammatory and anti-inflammatory cytokines. **(B)** Schematic of neuronal mechanism by which PDX inhibits TRPA1/TRPV1 and calcium signaling in nociceptor neurons via GPR37.

Our findings provided new mechanistic insights into macrophage signaling in pain. Mechanistically, PDX’s effect on macrophage phagocytosis and efferocytosis is highly dependent on intracellular calcium and requires the involvement of Giα, Gβ/γ, and the PI3K/AKT signaling pathway (Supplemental Figure 9A, B). Efferocytosis plays a crucial role in tissue homeostasis to ensure the removal of apoptotic cells (e.g., neutrophils) in tissues, preventing the accumulation of dead cells. SPMs have been shown to promote inflammation resolution by enhancing phagocytosis and efferocytosis (40, 41). Our findings show that PDX triggers GPR37-mediated phagocytosis in peritoneal macrophages via calcium signaling, where calcium chelator BAPTA-AM inhibits PDX-induced phagocytosis. Additionally, Gi signaling via GPR37 is necessary for phagocytosis, which completes within 60 minutes post-initiation, while efferocytosis of apoptotic neutrophils, linked to inflammation resolution, occurs hours later. Our bulk RNAseq results also demonstrate that PDX upregulates the S100 signaling pathway, enhancing genes associated with nitric oxide and ROS production in DRG macrophages, crucial for pathogen defense and macrophage migration, and influencing antiviral and antibacterial responses (Figure 6A-F). Consistent with our findings, increased levels of the neutrophil markers S100a9 and S100a8 have been linked to the resolution of inflammatory pain in mice, while neutrophil activation is also associated with recovery from low back pain in patients (6).

Growing evidence underscores the importance of macrophages in pain regulation. Macrophages promote pain through mediators like cytokines that interact with nociceptors (42–45), and their polarization under different conditions can both initiate and alleviate pain (46). Macrophages also facilitate the resolution of inflammatory pain and neuropathic pain through cytokine production (e.g., IL-10), phagocytosis, exosome secretion, or mitochondrial transfer (18, 47–49). While microglia predominantly regulate neuropathic pain in males (50), macrophages affect pain in both sexes, with studies revealing microglia-independent, macrophage-dependent peripheral sciatic pain in both male and female mice (51). Furthermore, the TLR9-mediated macrophage pathway regulates neuropathic pain in males (42), while the IL-23/IL-17-mediated macrophage signaling modulates mechanical pain in females (52). Our research indicates that PDX effectively reduces postoperative pain in both sexes following both early- and late-treatments (Supplemental Figure 5D, E). This is also reflected in the equal inclusion of male and female animals in most experiments. Sex and strain differences play an important role in pain processing (53, 54). We observed that female mice lacking GPR37 exhibit more severe and persistent cold pain compared to males, suggesting a chronic pain state driven by GPR37 deficiency in females. Additionally, we identified strain-specific differences in the resolution of postoperative pain: the duration of fPOP is shorter in CD1 mice than in C57BL/6 mice (2–3 weeks vs. 5–6 weeks). Heritability of nociceptive traits has been reported among inbred mouse strains, including C57BL/6J, across 12 nociceptive measures (54). CD1/ICR mice are outbred and generally exhibit more robust immune responses than inbred strains such as C57BL/6. Compared to C57BL/6 mice, immune cells from CD1 mice show heightened responsiveness to inflammatory stimuli (55, 56), which may contribute to accelerated fracture healing and shorter pain duration. CD1 mice also have larger body sizes and, in our hands, demonstrate greater resistance to LPS-induced septic death compared to C57BL/6 mice (57).

Furthermore, our studies have uncovered a neuronal signaling by PDX in alleviating acute pain and neurogenic inflammation (Figure 11B). It is established that SPMs modulate TRPA1 and TRPV1 activities in mouse DRG sensory neurons (9, 16). Our results showed that PDX did not inhibit AITC-induced calcium responses in TRPA1-expressing cells (Supplemental Figure 8D–E), indicating that PDX is not a direct TRPA1 antagonist. Instead, PDX may modulate TRPA1 and TRPV1 activity indirectly via GPR37 signaling in DRG neurons (Figure 11B), consistent with our previous reports showing that SPMs inhibit TRPA1 and TRPV1 through GPCR pathways (38, 58). Our RNA-seq data revealed that PDX treatment significantly reduced Trpa1 expression in the mouse DRG following bone fracture (Figure 10K), in support of our calcium imaging studies showing that PDX can significantly suppress bone fracture-induced calcium responses in DRG neurons ex vivo. To demonstrate translational relevance, our spatial transcriptomic analysis revealed co-localization of GPR37 with TRPA1 and TRPV1 in human DRG neurons, providing a cellular basis for GPR37-mediated regulation of these ion channels. By inhibiting both TRPA1 and TRPV1 pathways, PDX effectively reduced spontaneous pain, neurogenic inflammation, and calcium influx induced by AITC and capsaicin. Supporting this, pathway analysis and ex vivo calcium imaging demonstrated significant suppression of calcium signaling and reduced neuronal hyperactivity in DRG neurons following PDX treatment in the fPOP model (Figure 5J). Furthermore, in vivo electromyography confirmed that PDX attenuates C-fiber–evoked nociceptive muscle reflex responses. Future studies are warranted to investigate how GPR37 in sensory neurons regulates TRPA1 and TRPV1 signaling.

Notably, this research provides new insights into the resolution physiology mediated by SPMs (59) (Figure 5G). Our lipidomic analysis revealed that PDX is among the most abundantly produced SPMs in muscle and spleen tissues following orthopedic surgery (Figure 3A–F). Strikingly, PDX levels in muscle were approximately tenfold higher than those of PD1, underscoring its potential importance in inflammation and pain resolution. We previously demonstrated potent analgesic actions of RvD1 and RvE1 in animal models of inflammatory pain (9), and accordingly, we proposed the concept for the resolution of inflammation and pain by SPMs (8). Increasing evidence suggests anti-inflammatory treatments such as steroids and NSAIDs may delay the resolution of inflammatory pain (6, 8). Significantly, we found that treatment with dexamethasone and meloxicam delayed the resolution of fPOP, thereby extending pain recovery periods (Figure 5G). In contrast, PDX facilitated the resolution of fPOP by reducing pain intensity and shortening its duration. Additionally, our findings highlighted the pivotal role of GPR37 in resolving postoperative pain; in *Gpr37*^−/−^ mice, fPOP recovery was not observed within 3 months, indicating the development of chronic pain (60) (Figure 5G). Hence, our results support our initial hypothesis that unresolved acute pain can evolve into chronic pain (8). Our RNAseq experiments, pathway analysis, and micro-CT imaging indicate that PDX regulates macrophage/neutrophil signaling, tissue regeneration, and wound healing. Intriguingly, recent research revealed a critical role for neutrophil activation in resolving inflammatory pain (6). Consistently, we found marked immune activation in the DRG of PDX-treated mice in the fPOP (Figure 6A), as evidenced by the upregulation of the S100A family pathway, including *S100a8* and *S100a9* (Figure 6A-G). Correspondingly, PDX treatment increased IL-10 expression and secretion from pM𝜑 following LPS stimulation. As a key anti-inflammatory cytokine, IL-10 is critical for the resolution of pain (18).

Globally, over 310 million surgeries are performed each year, including 40 million orthopedic surgeries, with the U.S. accounting for 18.5 million in 2022 (61). Postoperative pain significantly affects recovery and quality of life, yet many patients report inadequate analgesia (62). Despite extensive research and advances in analgesic therapies, postoperative pain management remains insufficient (3, 31). A systematic review of major gastrointestinal surgeries found that preoperative use of ω-3 fatty acids may reduce hospital stay durations (63). However, these studies did not monitor the in vivo production of SPMs. In a large-scale ancillary study, middle-aged and older U.S. adults received moderate daily doses of omega-3 fatty acids did not experience reduction in the prevalence or severity of pain (64). A meta-analysis suggested that a higher daily dose of 2.7 g of omega-3s might be necessary to alleviate pain (65), yet these studies did not monitor in vivo SPM production. It is important to note that besides SPM production, these fatty acids undergo various metabolic processes in humans (66). Despite the limited efficacy of omega-3 fatty acids in pain management, our study demonstrates the superior analgesic potential of PDX via various administration routes (Supplemental Figure 7). Notably, mitigation of postoperative pain with extremely low intravenous doses of PDX (0.003 mg/kg ∼ 30 nM concentration) should not impact platelet function or increase bleeding risk. Furthermore, FDA-approved treatments such as the biased opioid agonist Olinvyk (TRV130, 0.1-0.5 mg/kg, i.v.) (67, 68) and VX-548 (60 mg, P.O.), a selective Na_V_1.8 sodium channel inhibitor, require much higher doses to alleviate postoperative pain (69).

This study has several limitations. While we identified GPR37 as a potential receptor for PDX, the involvement of other GPCRs in PDX-induced macrophage signaling cannot be ruled out. In sensory neurons, various GPCRs or ion channels could mediate PDX’s analgesic effects. Further research into the downstream signaling pathways of PDX in immune cells and neurons, is warranted. A recent study demonstrated that PD1 restores myogenesis, enhances muscle regeneration following injury, and improves muscle phenotype in a dystrophic mouse model (70). These findings raise the intriguing possibility that PDX may similarly promote muscle regeneration through GPR37 signaling.

In conclusion, we highlighted a “resolution physiology” (59) and “resolution pharmacology” (16) in this study. PDX is endogenously produced and exerts potent analgesic and pro-resolution effects through GPR37 activation and coordinated macrophage and neuronal signaling (Figure 11A–B). Notably, while anti-inflammatory treatments such as steroids and NSAIDs impair the resolution of fPOP, PDX promotes its resolution (Figure 5G). Given its enhanced potency, simpler production compared to PD1, and growing mechanistic understanding, PDX holds promise as a more effective and safer non-opioid therapeutic option for postoperative pain.

## Materials and Methods

### Animals

Adult mice (8-16 weeks) and young animals (4-5 weeks, for DRG calcium imaging only) were used in this study unless specifically described. *Gpr*37^tm1Dgen^ (JAX 005806) mice were purchased from the Jackson Laboratory (JAX). CD1 mice (Charles River Laboratories) were used for behavioral, electrophysiological, and biochemical tests. C57BL/6 mice were only used in experiments involving knockout mice. Animals were randomly assigned to each group. All animals were maintained at the Duke University Animal Facility. Sample sizes were based on our previous studies with similar assays (18). All animal experiments were approved by the Institutional Animal Care and Use Committees of Duke University.

### Sex as a biological variable

This study used both male and female animals.

### Lipidomic identification of PD1 and PDX

PD1 and PDX, used for *in vivo* and *in vitro* tests in this study, were purchased from Cayman Chemical and authenticated using UV spectroscopy and liquid chromatography/tandem mass spectrometry (LC-MS/MS), by comparing obtain UV spectra, retention times, and fragmentation patterns of the compounds with those of known standards.

To examine the production of SPMs in the muscle surrounding the tibia and the spleen following tibial bone fracture, muscle and spleen samples were collected 3 days after tibial bone fracture injury. Approximately 3 mg of muscle tissue from the fracture area and the entire spleen were immediately isolated and snap-frozen in liquid nitrogen. The tissue then underwent solid phase extraction (SPE) and resulting methanol fractions were analyzed using liquid chromatography tandem mass spectrometry, by matching retention times and fragmentation patterns to those of known standards. Details regarding LC-MS/MS equipment and conditions, mass spectrum, and the UV spectrum of PD1 and PDX are presented in Supplemental Figures 2 and 3 and Supplemental Tables 1 and 2.

### Mouse models of postoperative pain and acute inflammatory pain

Tibial fracture was performed under isoflurane anesthesia as we described previously (20). Muscles were dissociated following an incision on the left hind paw. A 0.38-mm stainless steel pin was inserted into the tibia intramedullary canal, followed by the osteotomy. The incision was sutured with non-absorbable silk suture (6–0). Acute inflammatory pain was induced by a single intraplanar injection of allyl isothiocyanate (AITC, 100 μg) or capsaicin (1 μg).

### Drug injection

All reagents were dissolved in sterile PBS and injected using a Hamilton microsyringe with a 30-gauge needle under a brief isoflurane anesthesia. The drug volumes were 10 μl for local intraplantar (i.pl.) and intrathecal (i.t.) injections, 20 μl for peri-sciatic nerve (p.SN.) injection, and 100 μl for intravenous (i.v.) and intraperitoneal (i.p.) injection. For oral gavage (p.o.), PDX (200 μl) was given by a reusable oral gavage needle (20G*50mm).

### Behavioral tests for evoked pain and spontaneous pain in mice

Mechanical pain was assessed using von Frey filaments. Animals were habituated to the testing environment daily for at least 2 days before baseline assessment. The room temperature and humidity remained stable for all experiments. To test mechanical sensitivity, we confined mice in boxes placed on an elevated metal mesh floor and stimulated their hind paws with a series of von Frey hairs with logarithmically increasing stiffness (0.02–2.56 g, Stoelting), presented perpendicularly to the central plantar surface. We determined the 50% paw withdrawal threshold by Dixon’s up-down method. Thermal sensitivity was tested using a Hargreaves radiant heat apparatus (IITC Life Science). The basal paw withdrawal latency was adjusted to 9 to 15 seconds, with a cutoff of 20 seconds to prevent tissue damage. To test cold sensitivity, a drop of acetone was applied to the plantar surface of a hind paw, and the mouse’s response was observed for 60 seconds after acetone application. Paw flicking and licking time (seconds) to acetone was recorded. For fracture-induced spontaneous pain, we measured the time (seconds) mice spent licking or flinching the affected hind paws over 2 minutes after the fracture surgery at multiple time points. For capsaicin and AITC-induced spontaneous pain, we measured the time (seconds) mice spent licking or flinching the affected hind paws over 5 minutes after the capsaicin injection (i.pl. 1 μg) or AITC (i.pl. 100 μg). Behavioral assessments were conducted in a blinded manner.

### Dot blot assay for lipid-protein binding

Lipid membrane coating and protein overlay assay were conducted as previously described (18). See more details in Supplemental Methods.

### ELISA

ELISA for IL-1β, TNF-α, and IL-10 was conducted according to the manufacturer’s instructions. See more details in Supplemental Methods.

### Quantitative real-time RT-PCR

Quantitative real-time PCR for *Il1b*, *Tnf*, was conducted as previously described (xxx). See more details in Supplemental Methods.

### Bulk RNA sequencing and pathway analysis of DRG tissues

Total RNA was extracted from L3–L5 DRG and used to prepare mRNA libraries with the RNeasy® Mini Kit (QIAGEN, Cat# 74104). All samples met quality criteria (RIN > 8.0, 260/280 > 2.0). Sequencing was performed on the Illumina NovaSeq 6000. Differential expression and pathway analysis were conducted to compare PDX- and vehicle-treated fracture groups. Pathway enrichment was assessed using gene ratio and Fisher’s exact test. Results were visualized with a volcano plot. Detailed protocols are provided in the Supplemental Methods.

### Phagocytosis and efferocytosis assay in pM𝜑s and THP1 cells

The phagocytosis assay was modified from a previously described protocol (18). pHrodo Red Zymosan Bioparticles (Thermo Fisher Scientific, catalog P35364) were rinsed and reconstituted in RPMI medium. Zymosan particles (20 μg/ml) were added to cultures of pM𝜑s or THP1 cells. After 0, 15, 30, 45, and 60 min of incubation with beads at 37°C, nonadherent beads were removed with cold PBS, and cells were fixed with 4% paraformaldehyde for 10 min. For neutrophil efferocytosis analysis, we collected whole blood (5 ml from 5 mice) to extract neutrophils. Mouse neutrophils were marked by CellTracker™ Red CMTPX Dye (1 Thermo Fisher Scientific, catalog C34552) and followed by UV light exposure for 10 min (250 mW/cm²) to induce neutrophil apoptosis. Then we added the apoptotic neutrophils (5,000 cells) to the cultured pM𝜑s or THP1 M𝜑s. At 0, 1, 2, and 3 h after the incubation at 37°C, nonadherent neutrophils were removed with cold PBS, and the remaining cells were fixed with 4% paraformaldehyde for 10 min. Three optical fields were photographed by Zess780 confocal microscopy for quantification of zymosan particles or neutrophils ingested by M𝜑s, conducted on 50 to 400 pM𝜑s or THP1 cells per condition or well, and triplicate were included for statistical analysis. FITC-conjugated anti-mouse F4/80 antibody (1:400, Sigma Aldrich MAPF152D) was used to mark pM𝜑s, and FITC-conjugated anti-human CD45 antibody (1:400, BioLegend 304054) was used to mark THP1 M𝜑s.

### Flow Cytometry Analysis of Efferocytosis in Bone Fracture Tissue

Wild-type (WT) and *Gpr37* knockout (KO) mice were subjected to bone fracture, and muscle tissue surrounding the fracture site was harvested 3 days post-injury. Data were analyzed using Cytobank Software (https://www.cytobank.org/cytobank). The gating strategy was illustrated in Supplemental Figure 14. See more details in Supplemental Methods.

### Spatial transcriptomic analysis in human DRG

Human dorsal root ganglia (DRG) were obtained from donors through the National Disease Research Interchange (NDRI) with exemption permission from the Duke University Institutional Review Board (IRB). Spatial transcriptomic analysis was conducted using the Visium HD Spatial Gene Expression Reagent Kits (CG000686). See more details in Supplemental Methods.

### Micro X-ray computed tomography

Micro x-ray computed tomography (Micro-CT) analyses were performed on the tibial bone from fracture mice using a Nikon XTH 225 ST scanner with the 225-kV source (Scanco, Southeastern, PA) as previously described (71). Micro-CT data was quantified to assess the fractured leg’s diameter and cross-sectional area as an indication of edema on day 3 post-fracture.

### Statistics

All data in the figures are expressed as the mean ± SEM. The sample size for each experiment was based on our previous studies involving such experiments (18). Statistical analyses were performed using GraphPad Prism 10.2 (GraphPad Software). Biochemical and behavioral data were analyzed using a two-tailed Student’s t-test (unpaired, 2-group comparisons), one-way ANOVA followed by Bonferroni’s or Tukey’s post hoc test, or two-way ANOVA, followed by Bonferroni’s or Tukey’s post hoc test. For the time course analyses, repeated measures of ANOVA were used, when applicable. The criterion for statistical significance was a *P* value of less than 0.05.

### Study Approval

The Institutional Animal Care & Use Committee of Duke University approved all the animal procedures. Animal experiments were conducted in accordance with the National Institutes of Health Guide for the Care and Use of Laboratory Animals.

### Data and Software Availability

No custom software was used in this study. RNAseq data were deposited with Geo accession number GSE309530.

## Supplemental Information

Supplemental information includes Supplemental Methods, 13 Supplemental Figures, 2 Supplemental Tables.

## ACKNOWLEDGMENTS

This study was supported by Duke University Anesthesiology Research Funds, NIH grants R01NS131812 and R61NS138215, and UG3NS143650 and DoD grants W81XWH2110885, W81XWH2110756, W81XWH-2210267, W81XWH-2210646 to R.R.J. S.B. was supported by NIH R03 grant DE032394. C.N.S. and lab are supported by NIH/NIHGMS R35GM139430.

## AUTHOR CONTRIBUTIONS

Y.L. and R.R.J. developed the project. Y.L., S.B., J.X., M.L., and S.C. conducted experiments and data analyses; R.R.J., C.N.S., and Y.L. participated in project development. J.J. conducted the mass spectrometry analysis of PD1, PDX, and SPMs under the guidance of C.N.S. Y.L. and R.R.J. wrote the manuscript, and other co-authors edited the manuscript.

## Supplemental Information

**Supplemental Figure 1.**
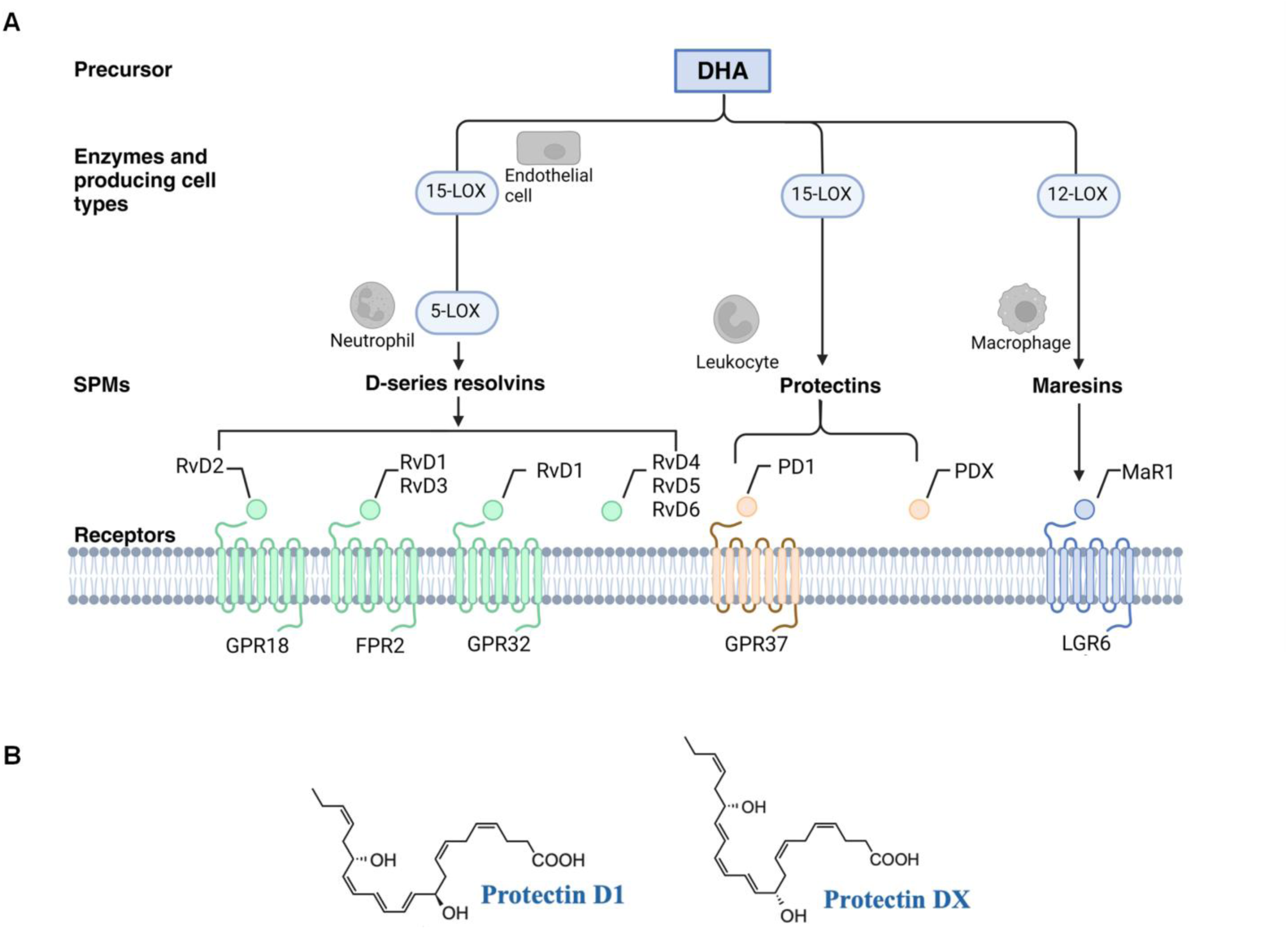
Biosynthesis of SPMs, NPD1, and PDX from omega-3 polyunsaturated fatty acids. (**A**) Schematic illustration of the pathway by which DHA (docosahexaenoic acid), a crucial precursor, is converted into specialized pro-resolving mediators (SPMs) through enzymatic reactions. The SPMs are classified into three main groups: D-series resolvins, protectins, and maresins. SPMs are produced by the combined actions of 15, 12, and 5 lipoxygenase (15-LOX, 12-LOX, 5-LOX) that interact with specific GPCRs, such as GPR18, FPR2, GPR32, GPR37, and LGR6, on immune cells, glial cells, and neurons. By binding to these receptors, SPMs help resolve inflammation and pain, facilitate tissue repair, and maintain homeostasis in the body (16, 41, 59, 72). (**B**) Structures of NPD1/PD1 (10R,17S-dihydroxy-4Z,7Z,11E,13E,15Z,19Z-DHA) and PDX (10(S),17(S)-dihydroxy-4Z,7Z,11E,13Z,15E,19Z-DHA).

**Supplemental Figure 2.**
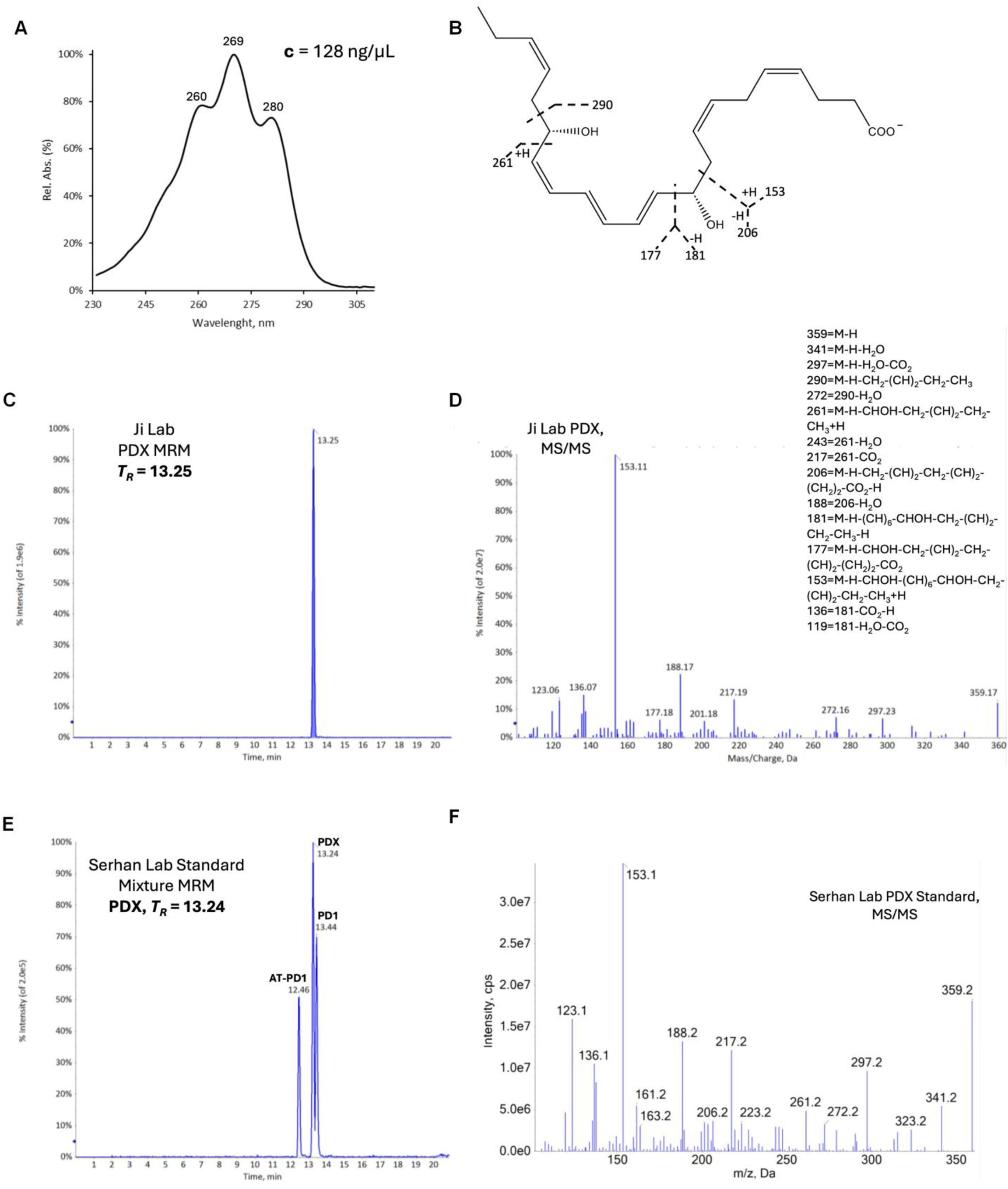
Authentication of PDX from Ji Lab (purchased from Cayman Chemical) in Serhan Lab. (**A**) UV absorbance spectrum of PDX at a concentration of 128 ng/μL (indicated as 100 ng/μL from Cayman). (**B**) structures of PD1. (**C-F**) Mass spectrometry (MS/MS) spectrum showing time (C, E) and mass/charge (D, F) to represent the fragmentation pattern of PDX from Ji lab (C, D) and Serhan lab (E, F) using Serhan Lab Standard.

**Supplemental Figure 3.**
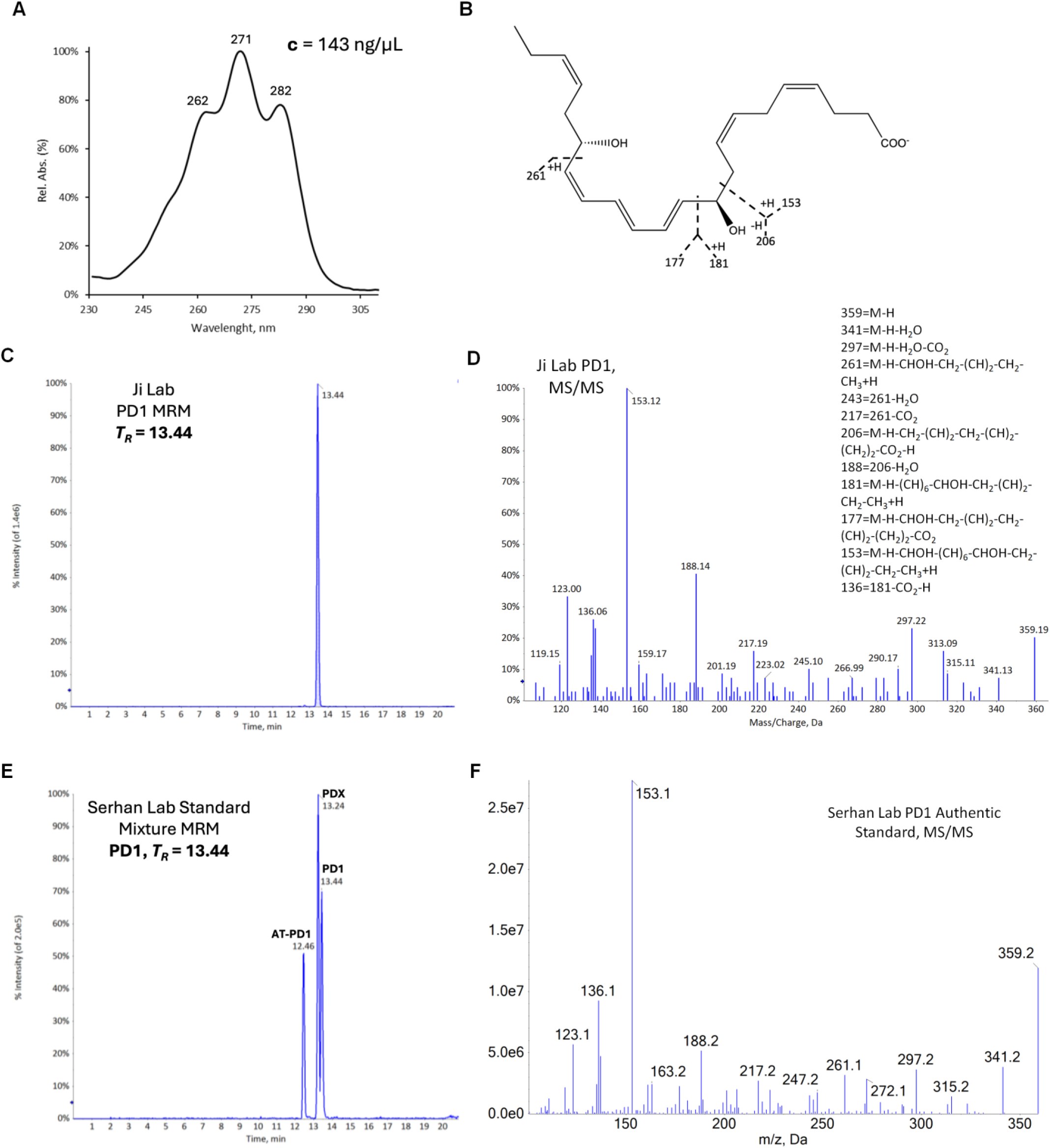
Authentication of PD1 from Ji Lab (purchased from Cayman Chemical) in Serhan Lab. (**A**) UV absorbance spectrum of PD1 at a concentration of 143 ng/μL (indicated as 100 ng/μL from Cayman). (**B**) structures of PD1. (**C-F**) Mass spectrometry (MS/MS) spectrum showing time (C, E) and mass/charge (D, F) to represent the fragmentation pattern of PD1 from Ji lab (C, D) and Serhan lab (E, F) using Serhan Lab Standard.

**Supplemental Figure 4.**
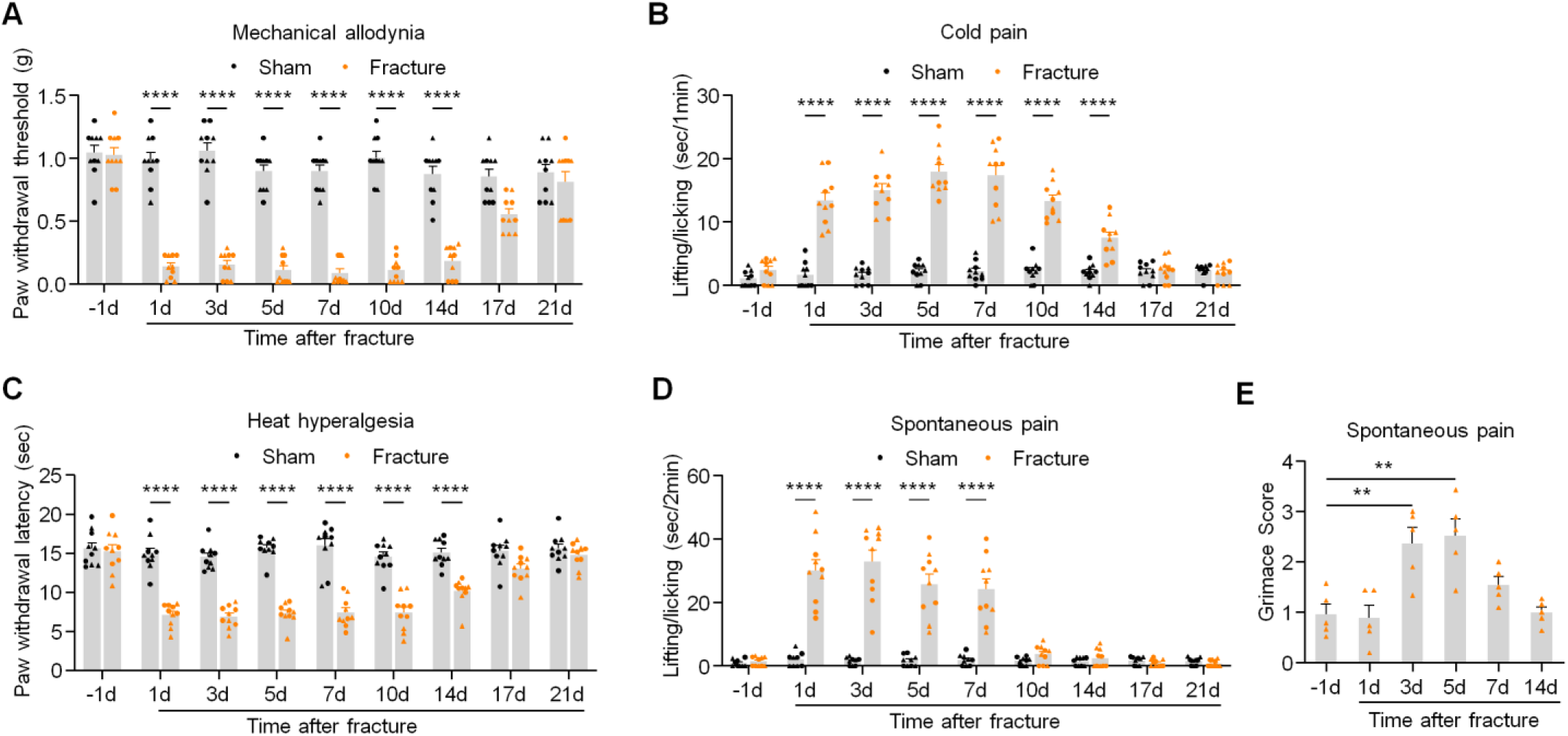
Time course of sham surgery and tibial fracture induced postoperative pain in CD1 mice. (**A**) Development and recovery of mechanical allodynia in von Frey test. (**B**) Development and recovery of cold allodynia in acetone test. (**C**) Development and recovery of heat hyperalgesia in Hargraves’ test. (**D**) Development and recovery of spontaneous pain as time spent on lifting/licking behavior. (**E**) Tibial fracture induced spontaneous pain evaluated by Grimace score. Data are represented as mean ± SEM and statistically analyzed by two-way ANOVA or one-way ANOVA with Bonferroni’s post hoc test. \*\**P* < 0.01, \*\*\*\**P* < 0.0001; *n* = 10 mice (5 males and 5 females, A-D), n = 5 males (E). ▴ male, ● female.

**Supplemental Figure 5.**
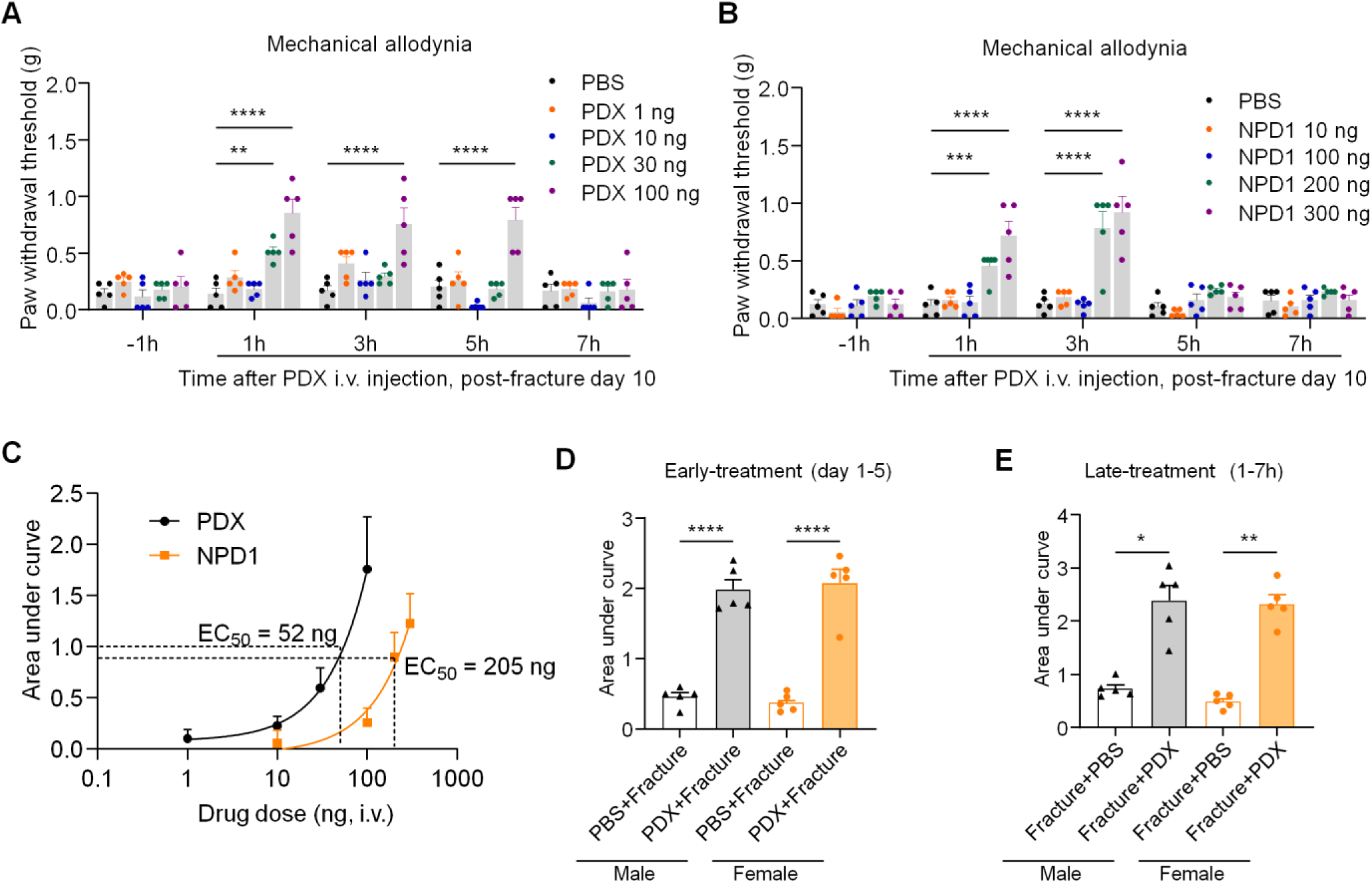
Dose-dependent effects of PDX and NPD1 in fPOP in CD1 mice. (**A**) Effects of i.v. injection of PDX (1, 10, 30, and 100 ng) and PBS (vehicle) on mechanical allodynia, revealed by PWT in von Frey tests on post-surgical day 10. n = 5 female mice per group. (**B**) Effects of i.v. injection of NPD1 (10, 10, 200, and 300 ng) and PBS on mechanical allodynia on post-surgical day 10. n = 5 female mice per group. (**C**) Comparison of AUC curves of PWT after PDX and NPD1 treatment, showing distinct EC_50_ values for PDX (52 ng) and NPD1 (205 ng). n = 5 female mice per group. (**D**) Comparison of AUC of PWT for PDX pre-treatment in males (black) and females (orange). Data were collected on post-surgical days 1, 3, and 5. n = 5 mice per sex per group. (**E**) Comparison of AUC of PDX post-treatment in males (black) and females (orange). Note that PDX is highly effective in reducing postoperative pain in both sexes. Data were collected from 1h, 3h, 5h, and 7h after PDX treatment on post-surgical day 10. n = 5 mice per sex per group. Data are represented as mean ± SEM and statistically analyzed by two-way ANOVA (A, B) and one-way ANOVA (C, D) with Bonferroni’s post hoc test. \**P* < 0.05, \*\**P* < 0.01, \*\*\**P* < 0.001, \*\*\*\**P* < 0.0001; ▴ male, ● female.

**Supplemental Figure 6.**
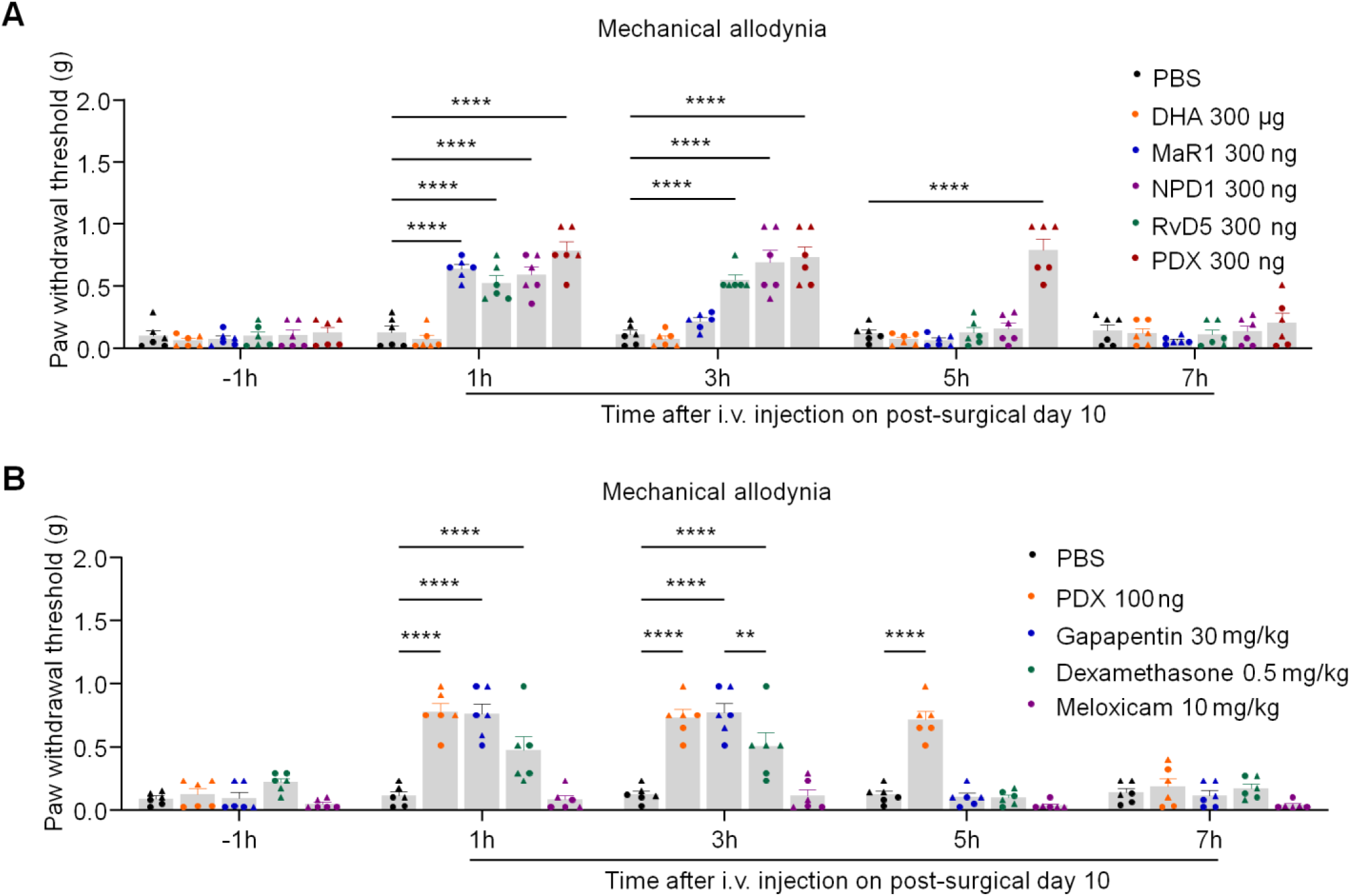
Comparison of PDX-induced analgesia with MaR1, NPD1, RvD5, DHA, dexamethasone, and gabapentin in fPOP in CD1 mice. (**A**) Effects of i.v. injection of DHA (300 μg), MaR1 (300 ng), NPD1 (300 ng), RvD5 (300 ng), and PDX (300 ng) on mechanical allodynia on fracture day 10. (**B**) Effects of i.v. injection of PDX (100 ng ∼3 μg/kg), gabapentin (30 mg/kg), dexamethasone (0.5 mg/kg), and meloxicam (10 mg/kg) on mechanical allodynia on fracture day 10. Note that PDX is much more potent than gabapentin, dexamethasone, and meloxicam in reducing postoperative pain. Data are represented as mean ± SEM and statistically analyzed by two-way ANOVA with Bonferroni’s post hoc test. \*\**P* < 0.01, \*\*\*\**P* < 0.0001; *n* = 6 mice per group (3 males and 3 females); ▴ male, ● female.

**Supplemental Figure 7.**
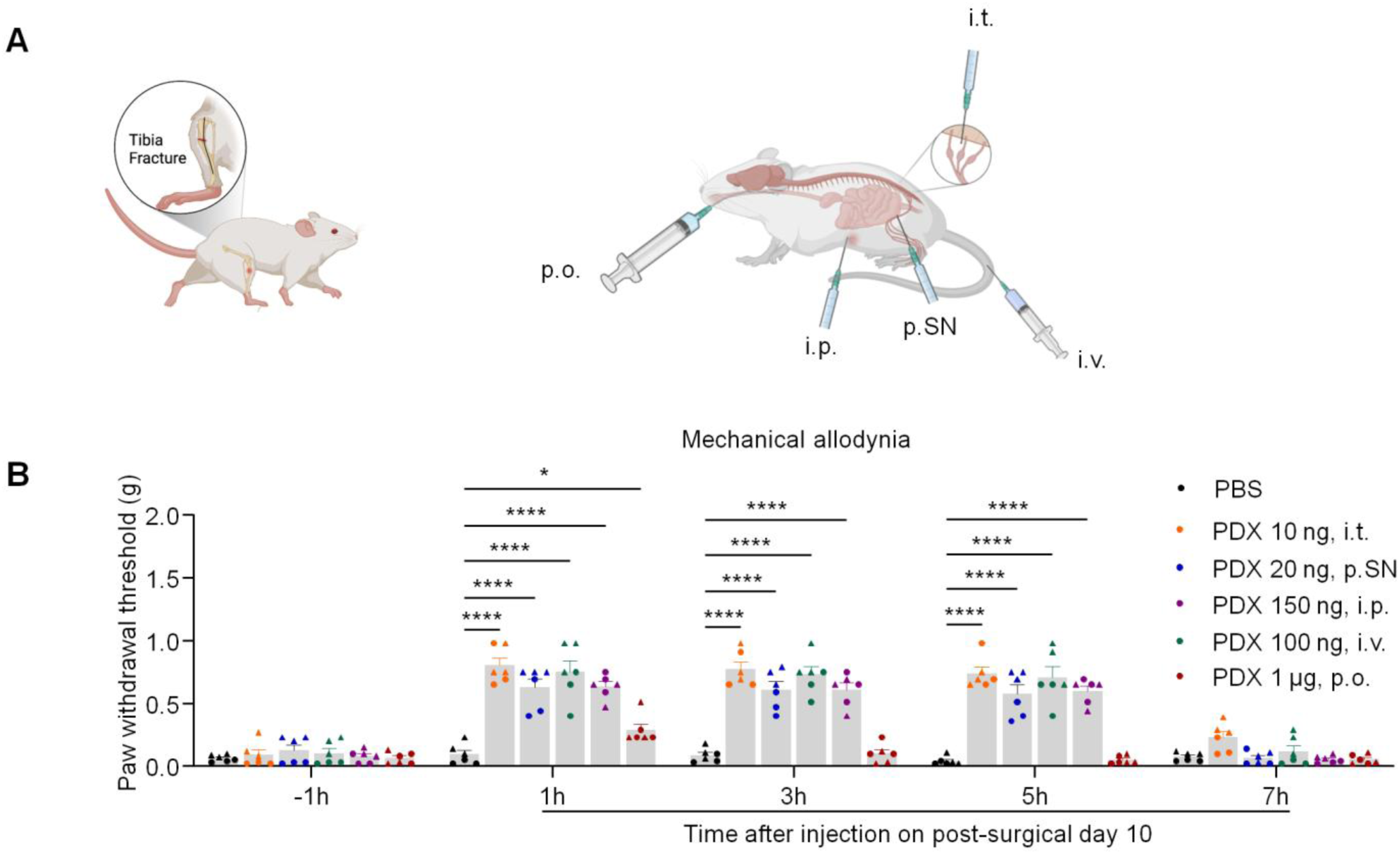
Comparison of PDX-induced analgesia in fPOP via different administration routes in CD1 mice. **(A)** Schematic of administration routes by which PDX inhibits postoperative pain. **(B)** Mechanical allodynia was assessed by PWT in von Frey tests, and PDX was administrated by intrathecal (i.t., 10 ng), peri-sciatic nerve (p.SN, 20 ng), intravenous (i.v., 100 ng), intraperitoneal (i.p., 150 ng), and oral gavage (p.o., 1 μg) routes on post-surgical day 10. Note that oral administration of PDX had only mild and transient pain relieving effects, even at a much higher dose. Data are represented as mean ± SEM and statistically analyzed by 2-way ANOVA with Bonferroni’s post hoc test. \**P* < 0.05, \*\*\*\**P* < 0.0001; *n* = 6 mice per group (3 males and 3 females); ▲ male, ● female.

**Supplemental Figure 8:**
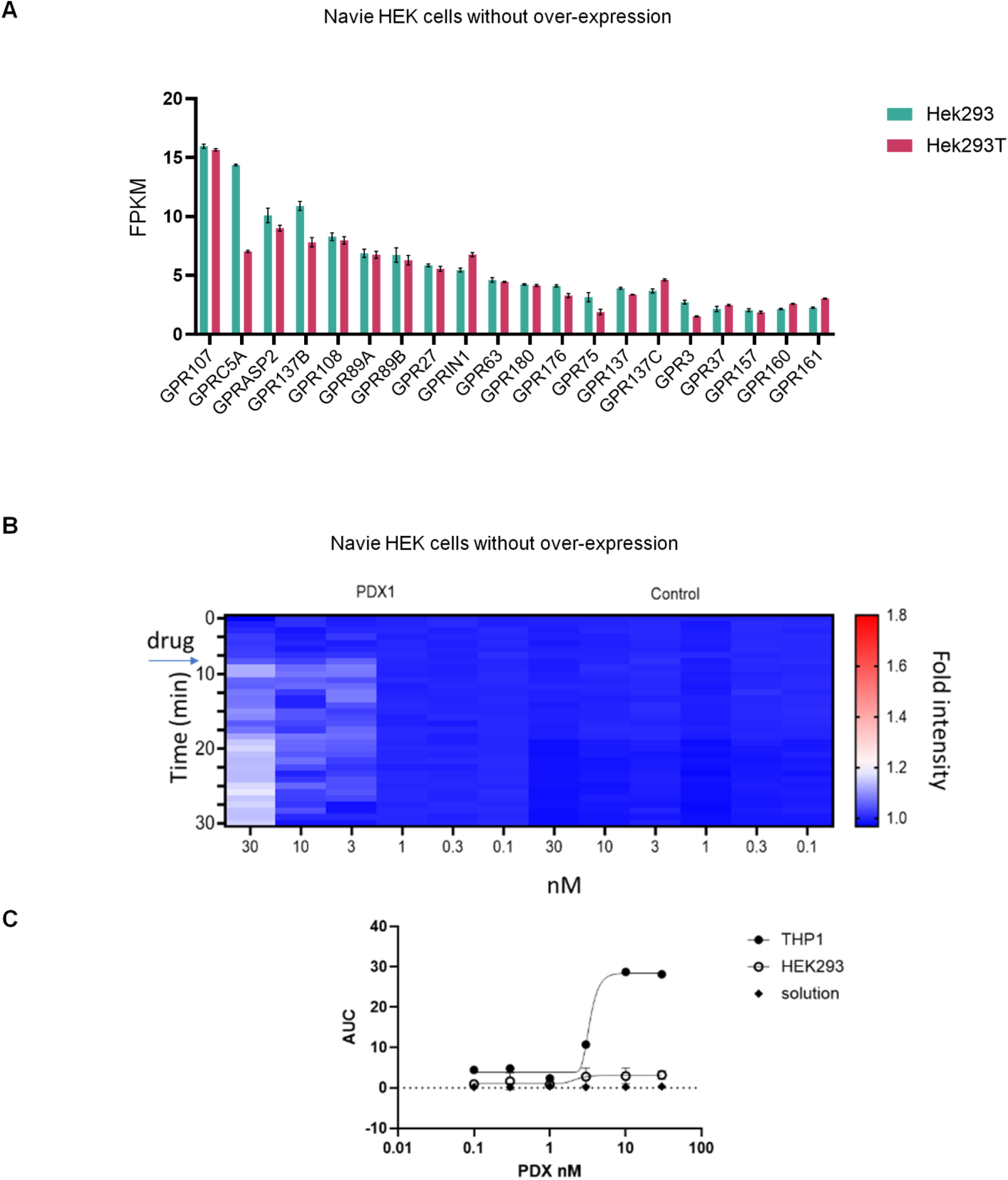
Characterization of basal constitutive activity of GPR37 expression in HEK293 cells. (A) Database analysis showing mRNA expression (FKPM value) of GPCRs in HEK293 and HEK293T cells (73). Notably, GPR37 expression is low compared to other GPCRs. (B) Native HEK cells exhibited no detectable response to PDX at low concentrations. Even at 30 nM, only ∼20% of cells showed a calcium response. (C) A dose–response experiment showing minimal Ca²⁺ signaling in naïve HEK cells. Notably, PDX induces robust Ca²⁺ response in THP1 cells, serving as a positive control (n =3).

**Supplemental Figure 9.**
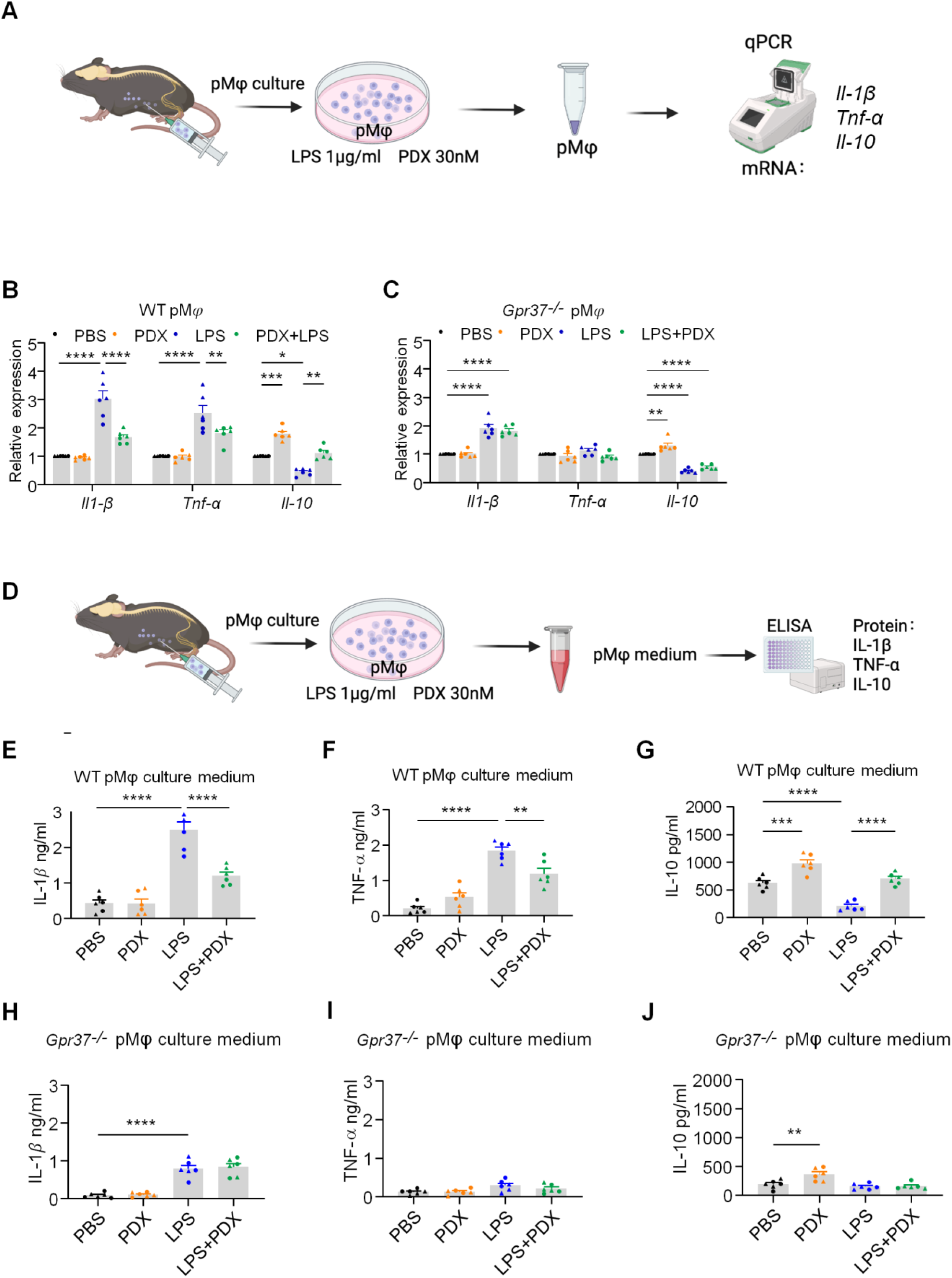
PDX regulates the expression of pro- and anti-inflammatory cytokines in peritoneal macrophages of C57BL/6 mice via GPR37. (A) Schematic of primary cultures for pM𝜑. Cultured cells were treated with LPS (1 μg/ml) and PDX (30 nM) for 24 hours and collected for qRT-PCR analysis for *Il1b*, *Tnf,* and *Il10* expression. (**B, C**) Relative mRNA expression levels of *Il1b*, *Tnf,* and *Il10* after LPS or PDX treatment in pM𝜑 cultures prepared from WT mice (B) and *Gpr37*^−/−^ mice (C). **(D)** Schematic of primary cultures for pM𝜑. Cultured cells were treated with LPS (1 μg/ml) and PDX (30 nM) for 24 hours, and the collected media to tested for ELISA IL-1β, TNF-α, and IL-10 expression. (E**-G**) Media protein levels of IL-1β (E), TNF-α (F), and IL-10(J) after LPS or PDX treatment in pM𝜑 cultures prepared from WT mice. (**H-J**) Media protein levels of IL-1β (H), TNF-α (I), and IL-10 (J) after LPS or PDX treatment in pM𝜑 cultures prepared from *Gpr37*^−/−^ mice. Data are represented as mean ± SEM and statistically analyzed by 2-way ANOVA with Bonferroni’s post hoc test (B, C) and 1-way ANOVA with Tukey’s post hoc test (E-J). \**P* < 0.05, \*\**P* < 0.01, \*\*\**P* < 0.001, \*\*\*\**P* < 0.0001; *n* = 6 cultures from 6 mice (3 males and 3 females); ▲ male, ● female.

**Supplemental Figure 10.**
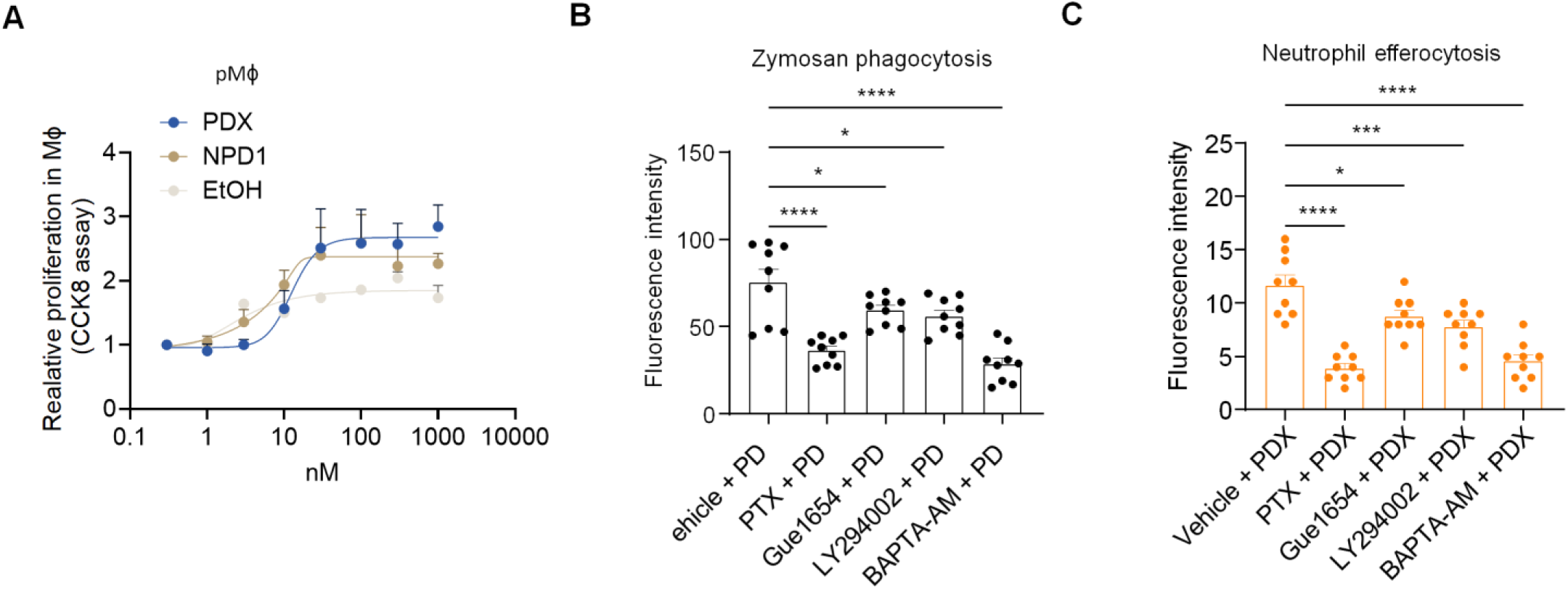
Effects of PDX on proliferation, phagocytosis, and efferocytosis of peritoneal macrophages of CD1 mice. **(A)** Dose-dependent effects of PDX and PD1 (0.3 nM to 1 µM) on macrophage proliferation activity in peritoneal macrophage culture using CCK8 assay. EtOH (0.3 nM to 1 µM) was used as a vehicle. Four cultures were prepared from 4 mice. EC_50_ = 12.5 nM for PDX. (**B, C**) Effects of PTX (1 μg/ml), Gue1654 (10 μM), LY294002 (20 μM), and BAPTA-AM (50 μM) on PDX (30 nM) induced zymosan phagocytosis (B) and neutrophil efferocytosis (C). *n* = 9 cultures/group. For each culture, the fluorescence intensity of zymosan and neutrophil uptake was analyzed by a plate reader. Data are represented as mean ± SEM and statistically analyzed by one-way ANOVA with Bonferroni’s post hoc test. \**P* < 0.05, \*\*\**P* < 0.001, \*\*\*\**P* < 0.0001.

**Supplemental Figure 11.**
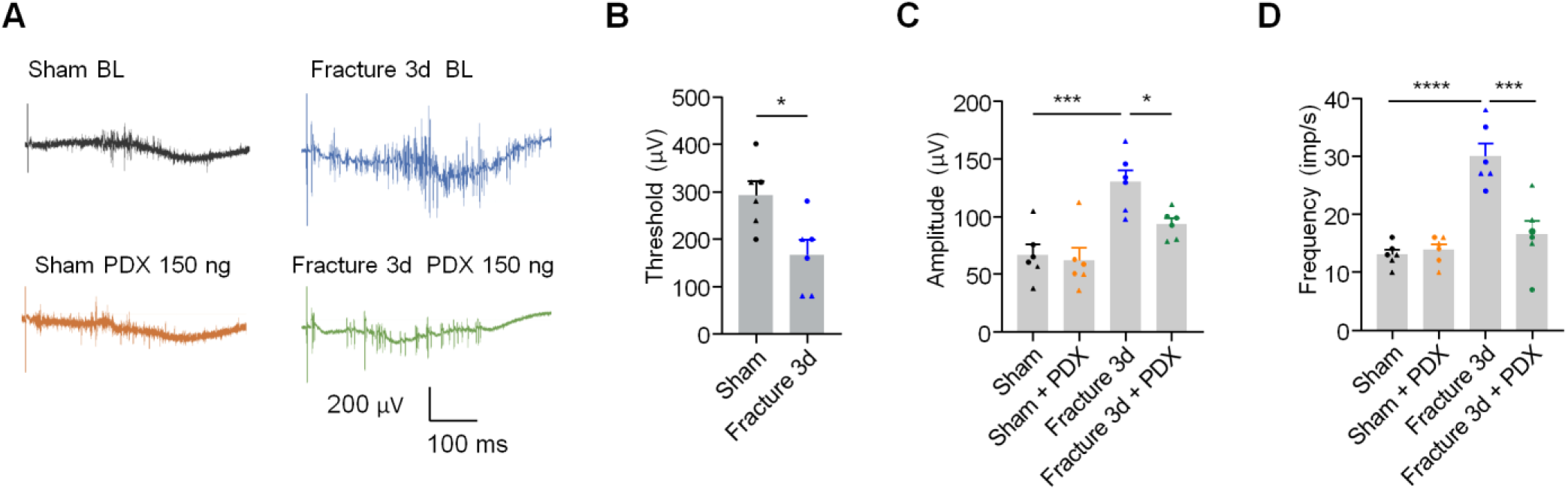
Intraperitoneal injection of PDX decreases the C-fiber reflex fPOP. **(A**) EMG traces of C-fiber reflex from 4 groups of animals with sham or fracture surgery with PBS and PDX. PDX (150 ng) was given by peritoneal injection. EMG recording was performed before (BL, baseline) and 30 min after the PDX treatment. (**B**) EMG threshold 3 days after sham and fracture surgery. (**C, D**) EMG amplitude (C) and frequency (D) from 4 groups in A. Data are represented as mean ± SEM and statistically analyzed by one-way ANOVA with Bonferroni’s post hoc test. \**P* < 0.05, \*\*\**P* < 0.001, \*\*\*\**P* < 0.0001; *n* = 6 mice per group (3 males and 3 females); ▴ male, ● female.

**Supplemental Figure 12.**
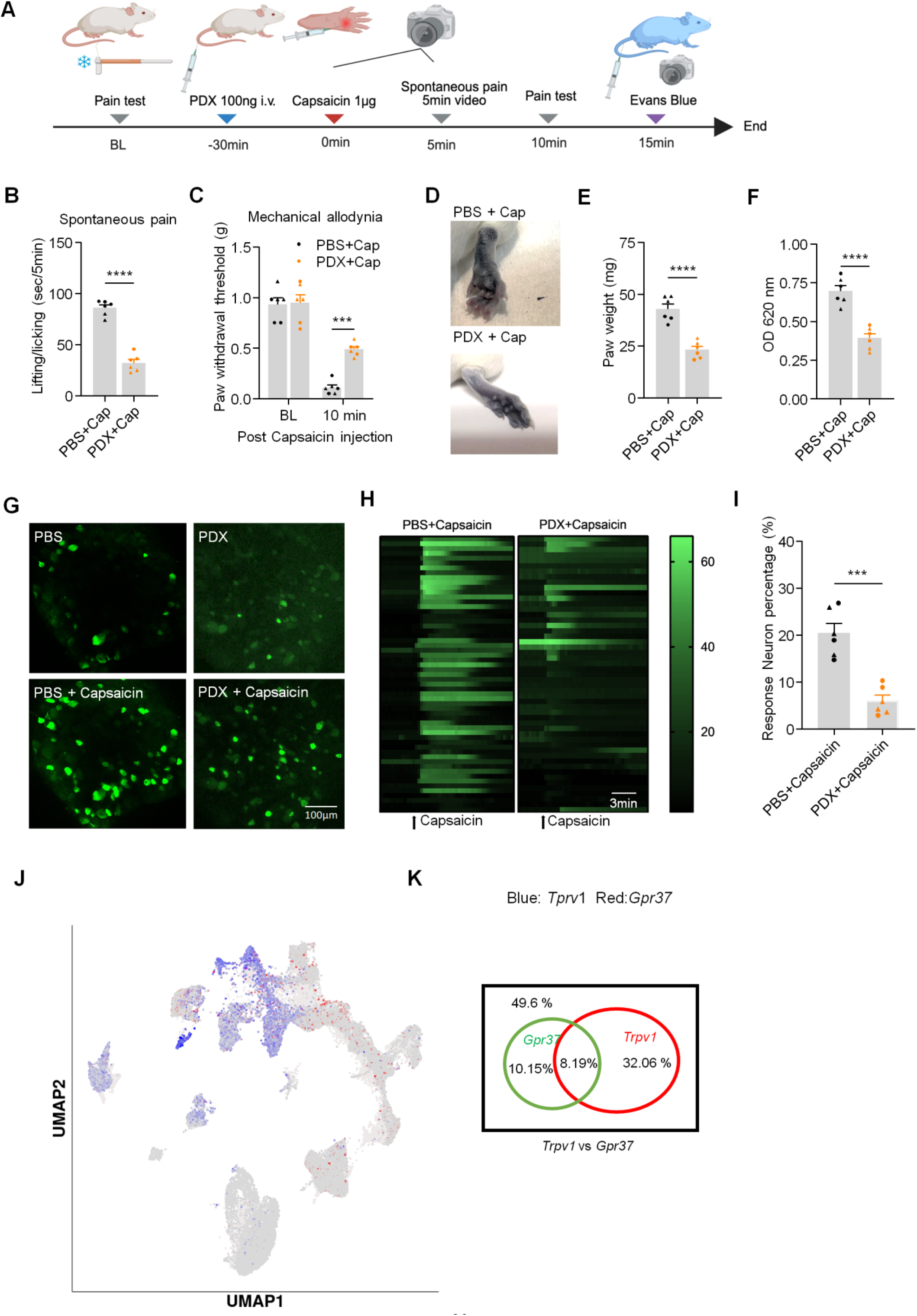
PDX reduces TRPV1-mediated spontaneous pain and neurogenic inflammation in CD1 mice. (**A**) Schematic of experimental design. Mice were treated with PDX for 30 min, followed by intraplantar injection of the TRPV1 agonist capsaicin (1 μg) and measurement of spontaneous pain. After the pain behavioral testing, Evans blue was injected intravenously to evaluate TRPV1-mediated neurogenic inflammation. (**B**) PDX blocks capsaicin-induced spontaneous pain, shown as reduced paw lifting or licking time within 5 min. (**C**) PDX blocks capsaicin-induced mechanical allodynia by von Frey test 10 min after injection. (**D**) Hind paw images showing capsaicin-induced edema 15 min after the Evans blue injection. (**E, F**) Quantification of paw edema by weight (E) and Evans blue concentration at OD 620 nM (F) in hind paws of mice treated with capsaicin with or without PDX (100 ng, i.v.). (**G**) DRG calcium images from four groups treated with PBS, PDX (30 nM), PBS + capsaicin (300 nM), and capsaicin + PDX. Scale bar, 100 μm. Note that Figure S12G is the same image as Figure 10H. (**H**) Heat map of DRG calcium signal in PBS + capsaicin and PDX + capsaicin groups. Each group consists of 51 DRG neurons. Scale bars, 3 min. (**I**) Percentage of capsaicin-responsive DRG neurons showing the effects of PDX pretreatment. (**J, K**) Co-localization of *Gpr37* with *Trpv1* in mouse DRG based on single-cell RNA-seq data. (J) UMAP plot showing *Gpr37* (blue) and *Trpv1* (red) expression. (K) Percentage of *Gpr37*/*Trpv1* co-expressing cells in mouse DRGs. Data are represented as mean ± SEM and statistically analyzed by unpaired Student’s t-test (B, E, F, I) and one-way ANOVA with Bonferroni’s post hoc test (C). \*\*\**P* < 0.001, \*\*\*\**P* < 0.0001; *n* = 6 mice per group (3 males and 3 females), ▲ male, ● female.

**Supplemental Figure 13.**
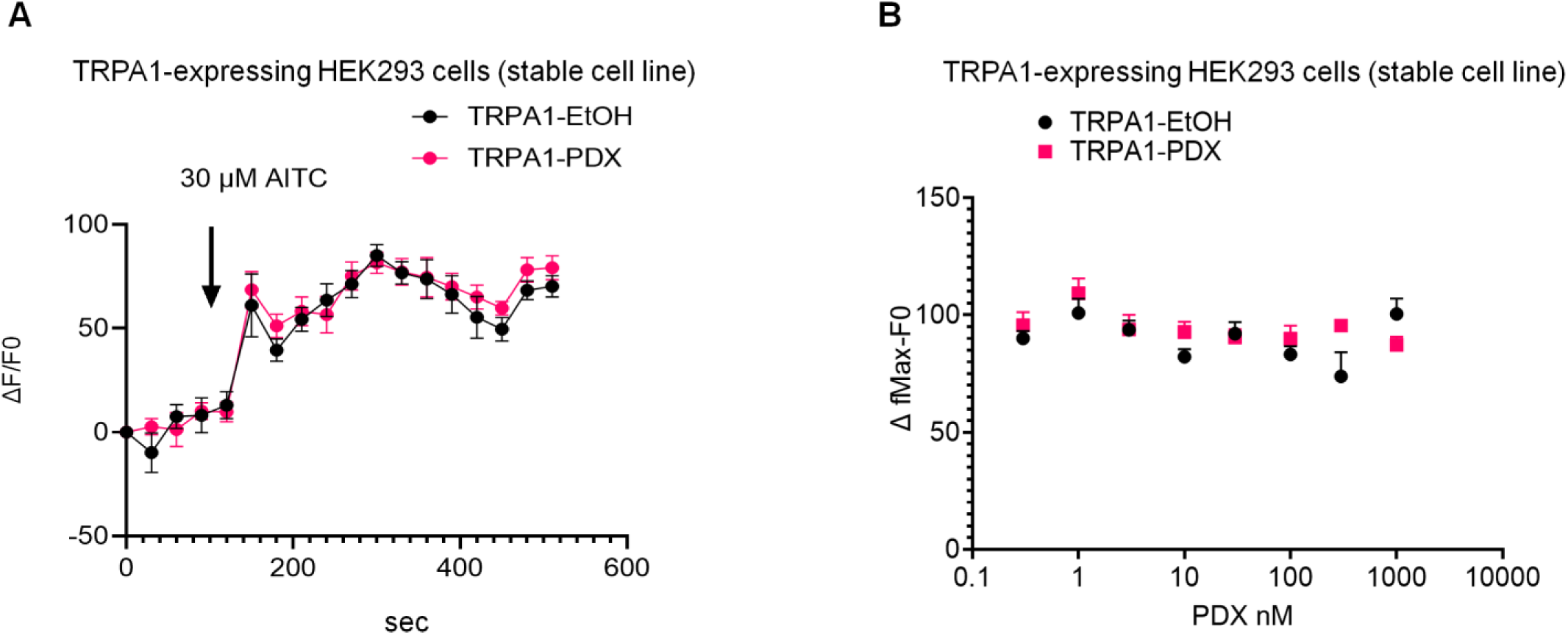
PDX has no direct effects on TRPA1 activity in TRPA1-expressing HEK293 cells. (**A**) Effects of 30 nM PDX on AITC-induced calcium influx in TRPA1-expressing HEK293 cells, visualized using Fluo-8 calcium imaging (n = 6 wells). (**B**) Dose–response curve of PDX effects (1 nM – 300 nM) on AITC (30 µM)-induced calcium influx in TRPA1-expressing HEK293 cells. Ethanol (EtOH) was used as vehicle control (n = 6 wells per condition).

**Supplemental Figure 14.**
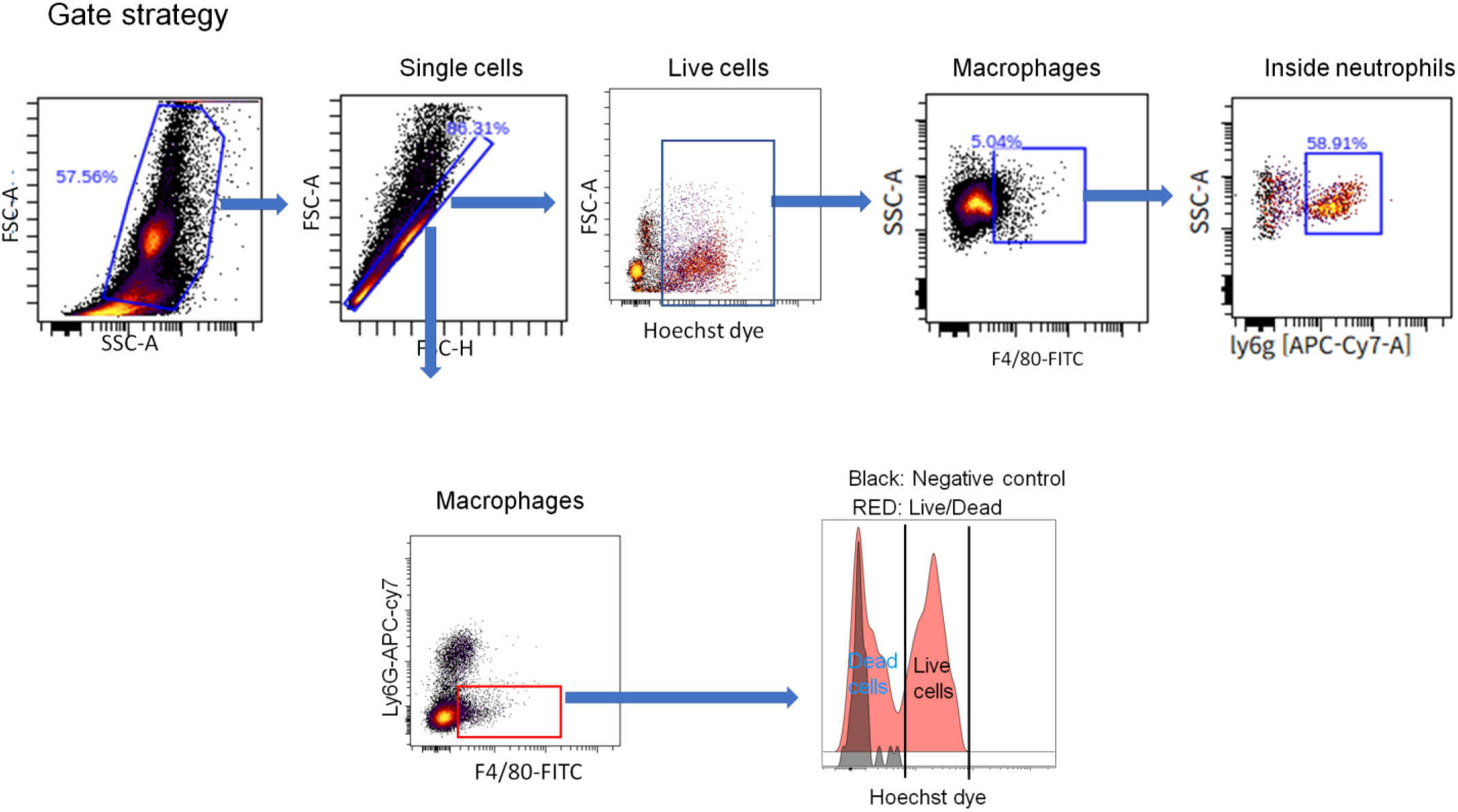
Gating strategy for in vivo efferocytosis detection in muscles of bone fracture area using flow cytometry.

**Supplemental Table 1:**
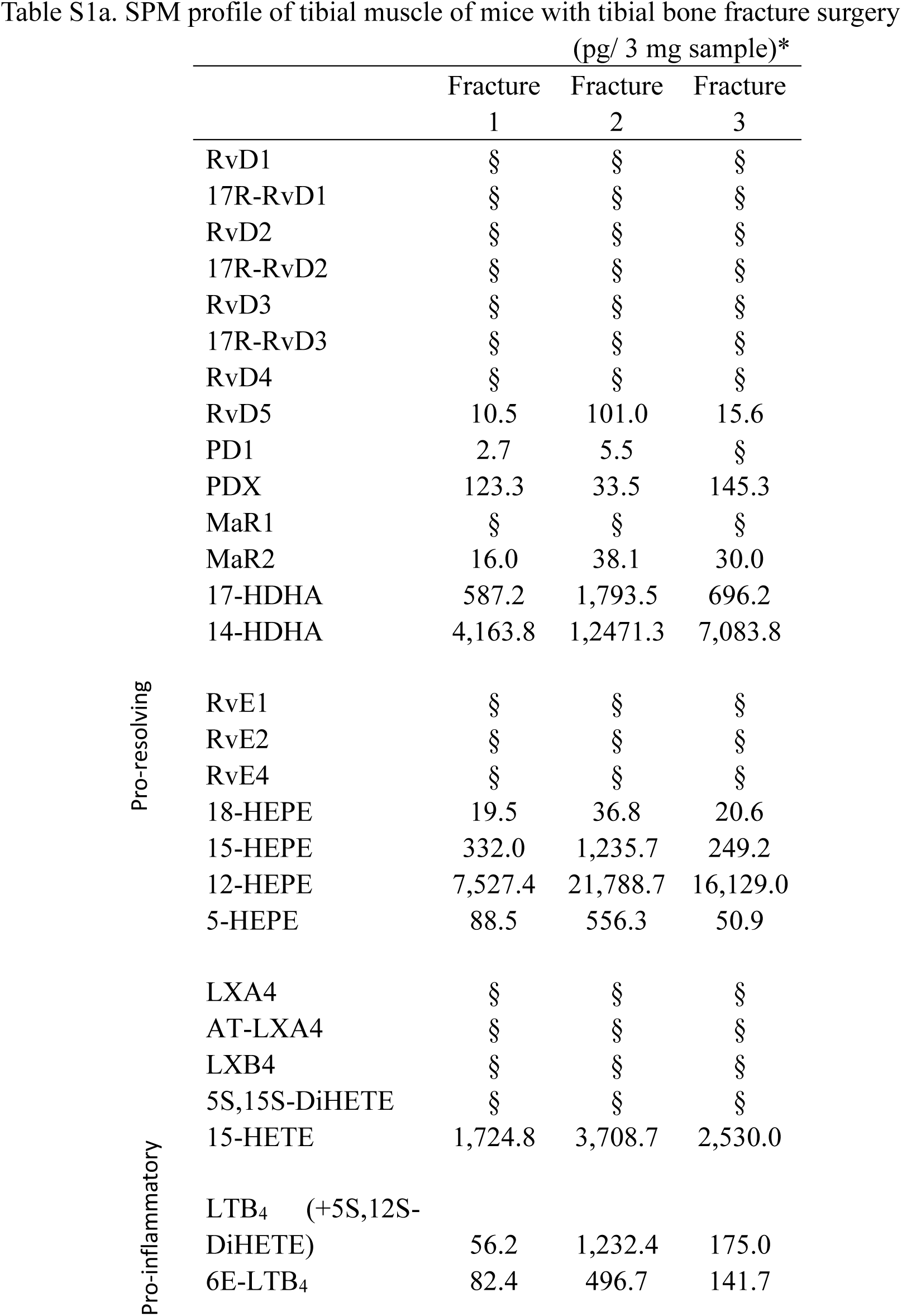
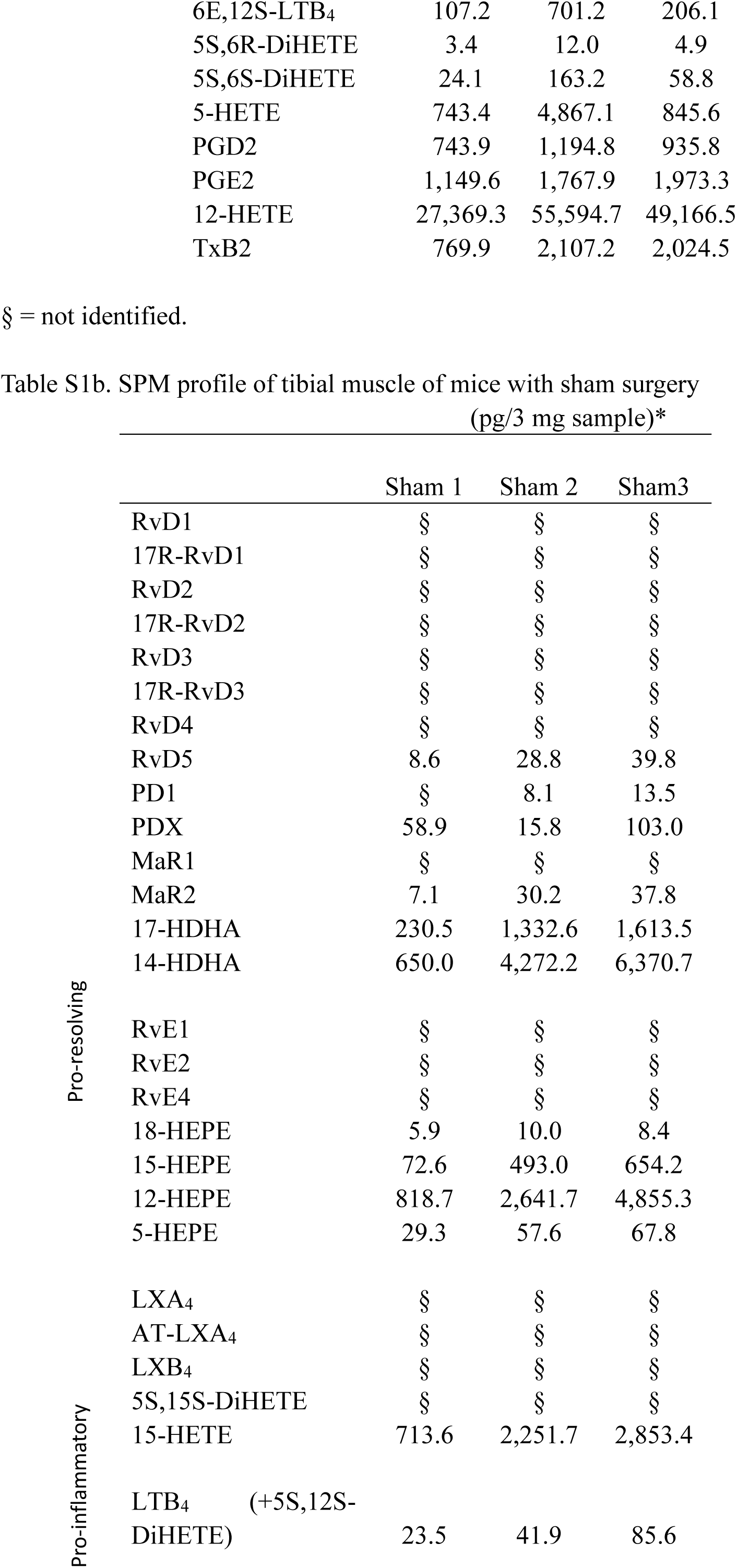
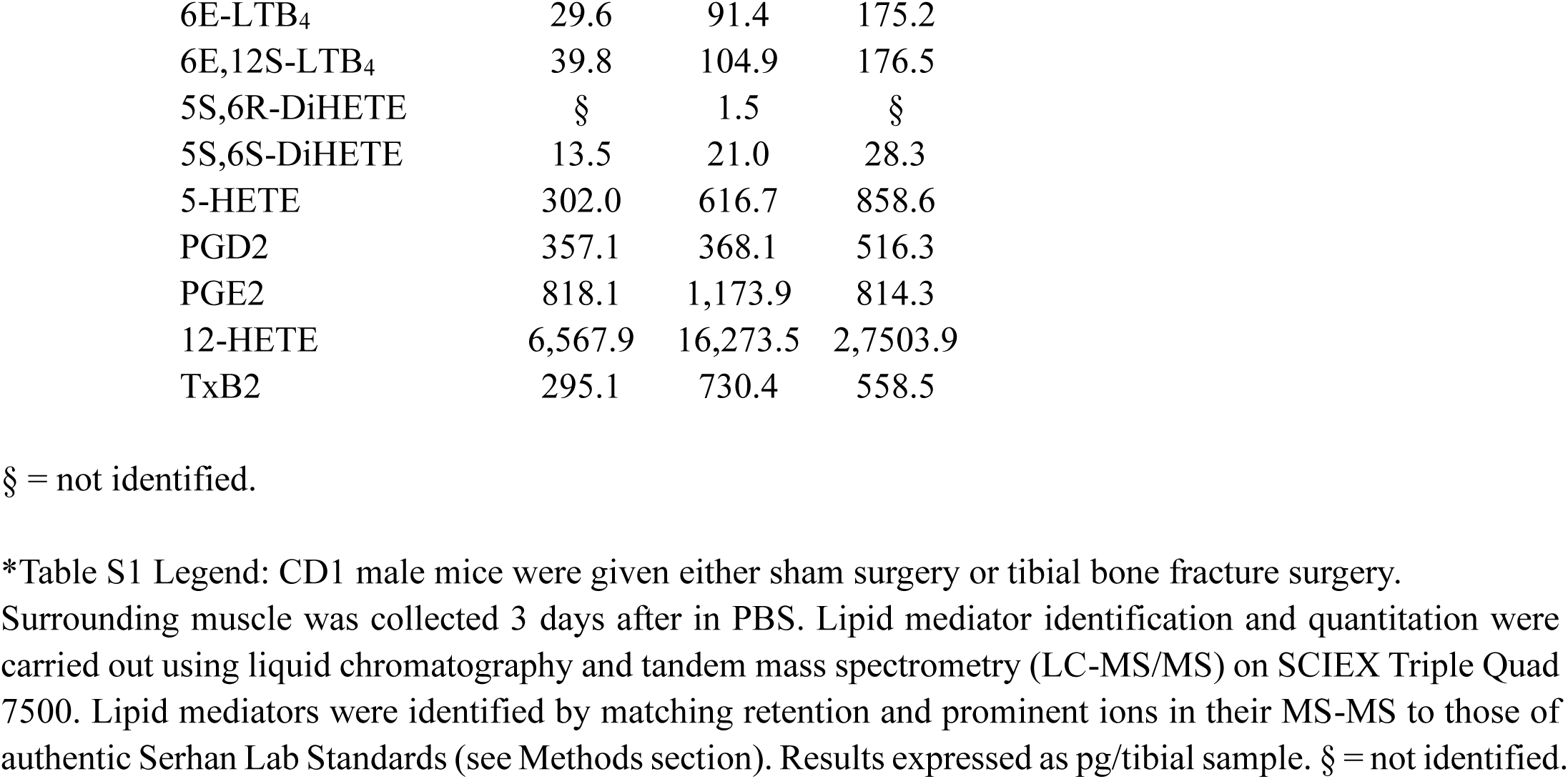
Lipidomic analysis in muscle near bone fracture of CD1 mice. Mice were given either sham surgery or tibial bone fracture surgery, and muscle was collected 3 days after in PBS. Lipid mediator quantification was carried out using liquid chromatography and tandem mass spectrometry (LC-MS/MS) on SCIEX Triple Quad 7500. Lipid mediators were identified by matching retention time and prominent ions in their MS-MS to those of authentic Serhan Lab Standards (see Methods section). n = 3 male mice per group. Results expressed as pg per muscle sample (3 mg tissue per muscle sample). § = not identified. (See below)

**Supplemental Table 2:**
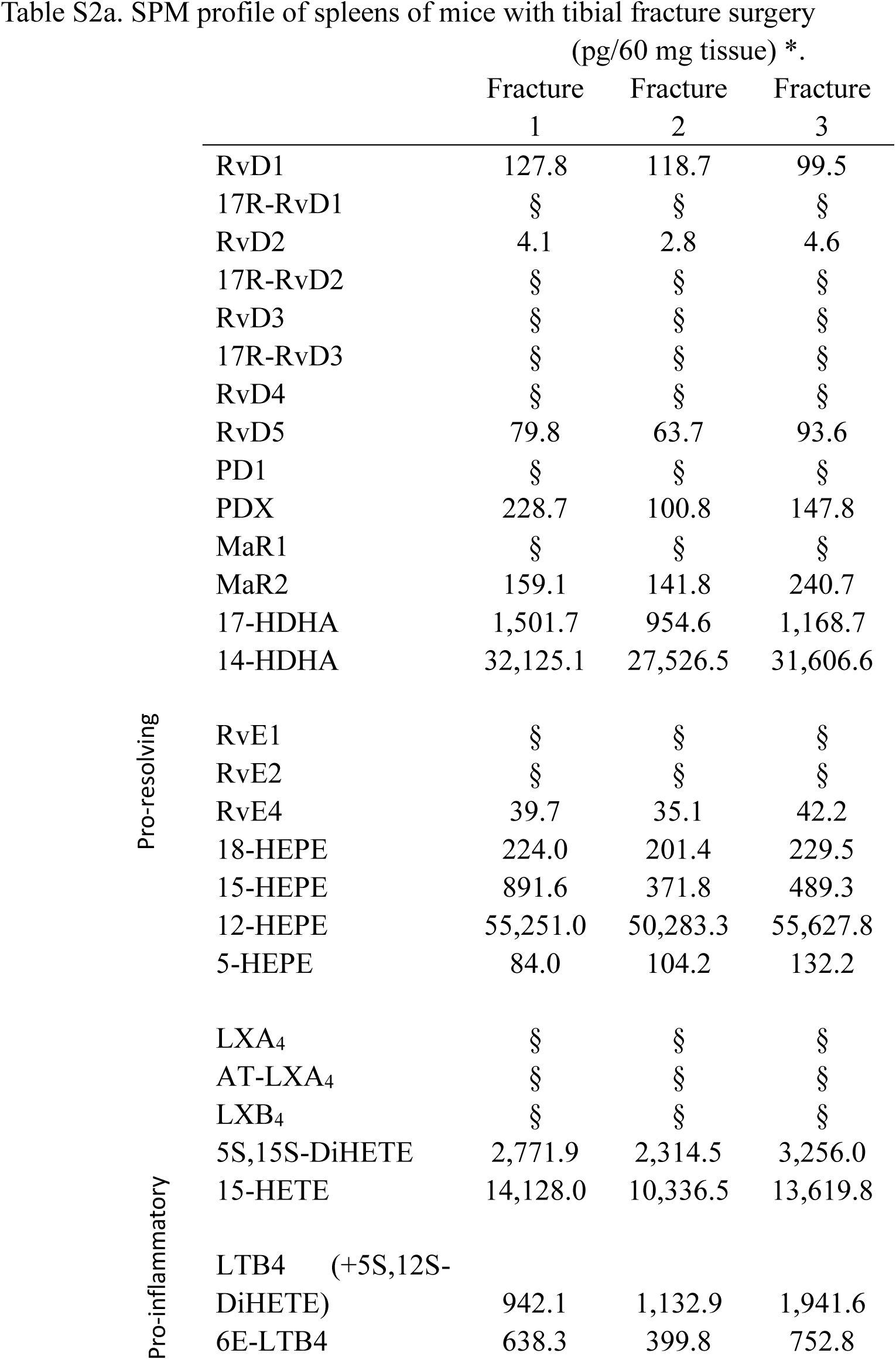
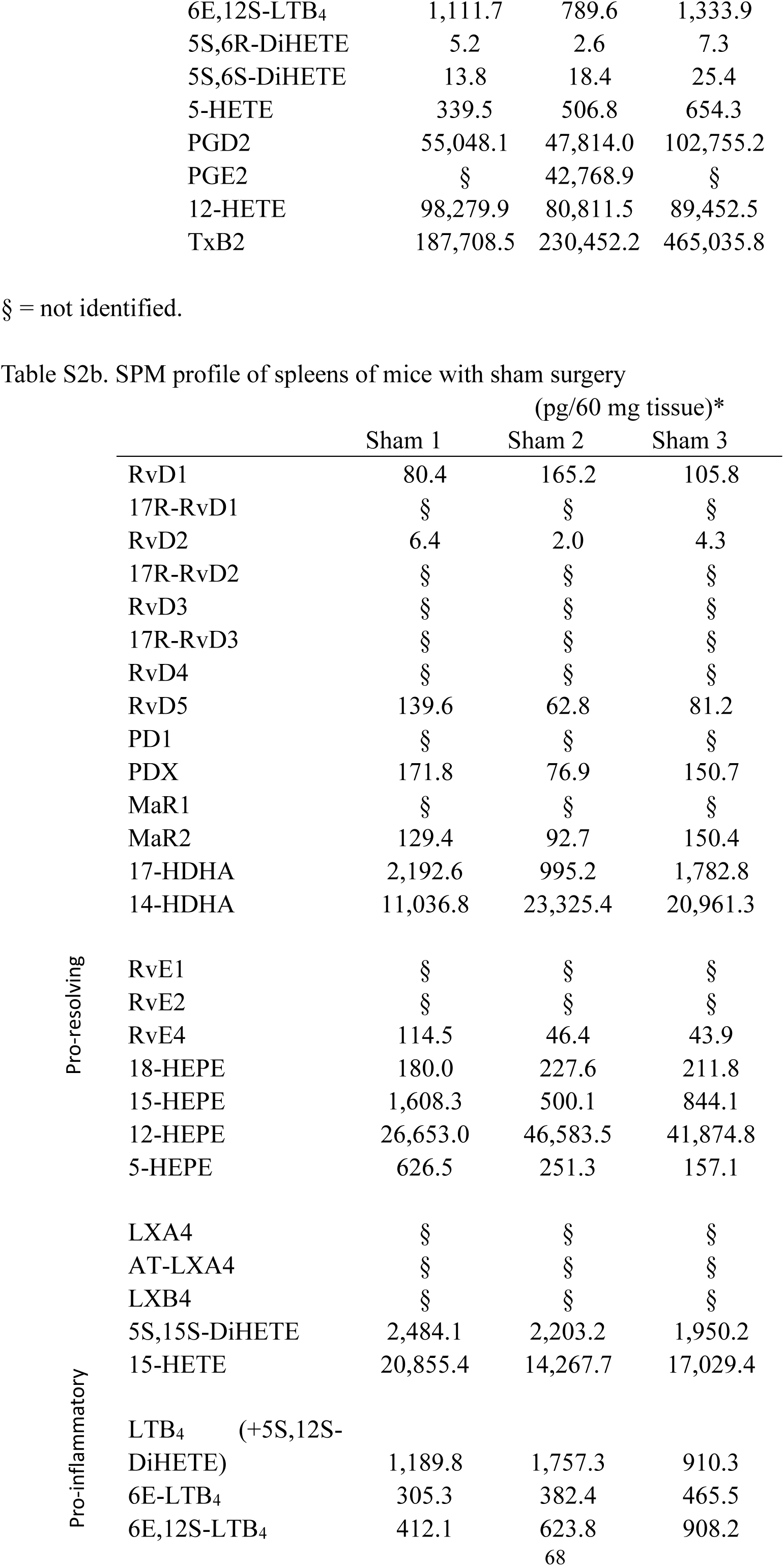
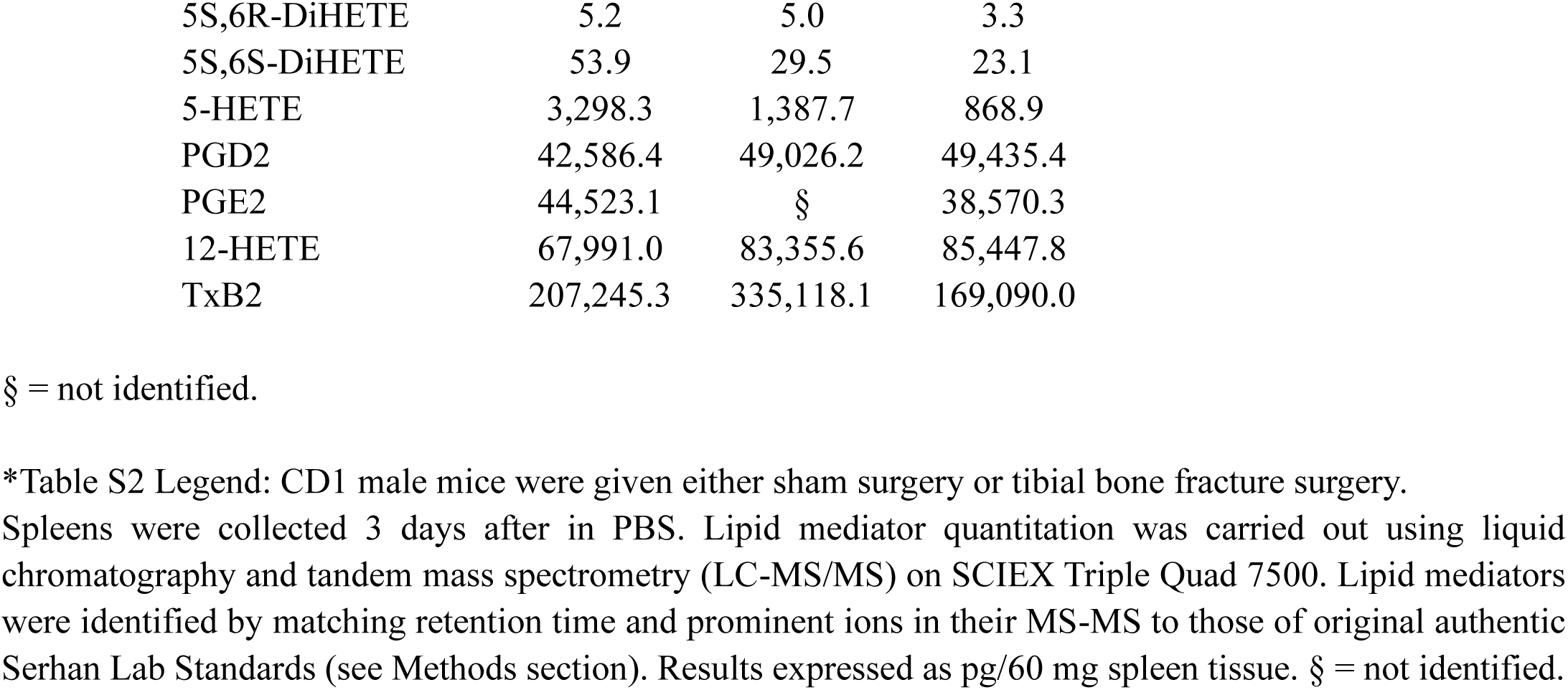
Lipidomic analysis in the spleen of CD1 mice. Mice were given either sham surgery or tibial bone fracture surgery, and the spleen was collected 3 days after in PBS. Lipid mediator quantification was carried out using liquid chromatography and tandem mass spectrometry (LC-MS/MS) on SCIEX Triple Quad 7500. Lipid mediators were identified by matching retention time and prominent ions in their MS-MS to those of authentic Serhan Lab Standards (see Methods section). n = 3 male mice per group. Results expressed as pg/60 mg spleen tissue. § = not identified. (See below)

## Supplemental Materials and Methods

### Cell cultures of HEK293T cells, DRG cells, pM𝜑s, and THP1 cells

#### HEK293T cell culture and transfection

The HEK293T flip-in cell line was purchased from the Duke Cell Culture Facility. The HEK293-hTRPA1 stable cell line was purchased from SB Drug Discovery (Cat # SB-HEK-TRPA1). Cells were cultured in Dulbecco’s Modified Eagle’s Medium containing 10% (v/v) FBS (Gibco, Thermo Fisher Scientific). Transfection (2 μg cDNA) was performed with Lonza electroporation at 70% confluency, and the transfected cells were cultured in the same growth medium for 48 h before use. The hGPR37-V5 pLenti304 plasmids were obtained from the DNASU Plasmid Repository.

#### Primary pM𝜑 culture

pM𝜑s were collected by peritoneal lavage with 5 ml warm PBS containing 1 mM EDTA. Cells were incubated in DMEM supplemented with 10% FBS at 37°C for 1 hour in a Petri dish and washed with PBS to eliminate nonadherent cells. The adherent cells were used as pM𝜑s. pM𝜑s were used after 2-3 days of culture.

#### THP1 cell culture

THP1 cell line was purchased from the Duke Cell Culture Facility. Cells were cultured in RPMI-1640 Medium containing 10% (v/v) FBS (Gibco, Thermo Fisher Scientific) and 0.05 mM 2-mercaptoethanol. To induce THP1 monocyte differentiation to macrophages, 25 nM of phorbol 12-myristate 13-acetate (PMA) was added to cultures.

### *In vitro* calcium imaging in HEK293T cells and pM𝜑s

HEK293T cells were loaded with 5 μM Fura2-AM (Invitrogen, Thermo Fisher Scientific, F1221) for 45 min and then resuspended in normal external buffer (140 mM NaCl, 5 mM KCl, 2 mM CaCl2, 2 mM MgCl2, 10 mM HEPES, titrated to pH 7.4 with NaOH) or external Ca^2+^-free buffer (140 mM NaCl, 2 mM MgCl2, 5 mM EGTA, 10 mM HEPES, titrated to pH 7.4 with NaOH). pM𝜑s collected from WT and *Gpr37*^−/−^ mice and TRPA1 HEK293 cells were loaded with 3 μM Fluo8-AM (AAT Bioquest, 21055) for 30 min and then resuspended in DMEM medium. Images of HEK293T cells with an excitation wavelength of 340 nm and 380 nm or pM𝜑s with an excitation wavelength of 488 nm were captured with a cooled Digital CMOS camera (ORCA-Flash 4.0, Hamamatsu Photonics). The ratio of fluorescence intensity of the two wavelengths in each experiment, as the peak amplitude/intensity in the first 10 min after treatment, was analyzed using MetaFluor software (Molecular Devices). The Shutter speed and wavelength were controlled by the pe-300 Fura system (Cool LED). Values from each experiment were normalized to the baseline ratio of 340:380 nm or the basal intensity for pM𝜑s.

### *Ex vivo* calcium imaging in whole-mount DRG from AAV-MaCPNS.2-hSyn-Gcamp6f-infected mice

AAV-MaCPNS.2-hSyn-Gcamp6f was intraperitoneally (i.p.) administered to C57BL6 mice (3×10^11^ vg /mouse) on postnatal day 1 (P1). Four weeks after the AAV injection, L3-L5 DRG were collected for an ex vivo DRG calcium imaging experiment. Mice were anesthetized through i.p. injection of 1.5 g/kg urethane, and their body temperature was maintained at approximately 37°C using a heat pad throughout the procedure. DRG were harvested and kept in ACSF. Stable imaging was achieved by placing a mash on the DRG to fix the position of DRG. Live imaging was facilitated using a Zeiss 780 upright confocal microscope with a 20X water objective lens, with the focal plane depth ranging from 50 to 70 μm. Seven images were captured per cycle, and 50 cycles were recorded. Following the acquisition of baseline measurements for 3 min, drugs (diluted in ACSF) were added to dishes, and a total of 15 min or 25 min time-lapse recordings were performed for each DRG. The data were analyzed using FIJI software.

### Dot blot assay for lipid-protein binding

Lipid membrane coating and protein overlay assay were conducted as previously described (18). Ethanol and chloroform-soluble fatty acids were directly loaded onto a hydrophobic PVDF membrane (MilliporeSigma). The membrane was coated with PDX and NPD1. The fatty acid–coated membranes were dried and blocked with 1% BSA. To express and isolate GPR37, HEK293 cells were transfected with GPR37 cDNA with a V5 tag (GPR37-V5) or with an empty vector (Mock transfection) using Lipofectamine 3000 (Invitrogen, Thermo Fisher Scientific). Cell lysates were incubated overnight with the coating membrane at 4°C, and the binding was detected by anti-V5–tagged antibody (mouse, 1:1,000, Thermo Fisher Scientific, catalog 46-0705). The blots were further incubated with an HRP-conjugated secondary antibody and developed in ECL solution (Pierce, Thermo Fisher Scientific). The intensity of lipid-protein binding was evaluated using ImageJ software.

### ELISA

Mouse ELISA kits for IL-1β, TNF-α, and IL-10 were purchased from R&D Systems (catalog MLB00C for IL-1β, catalog MTA00B-1 for TNF-α, and catalog M1000B for IL-10). ELISA was performed using culture media of pM𝜑s. For each ELISA assay, 50 μl of culture medium was used. ELISA was conducted according to the manufacturer’s instructions. The standard curve was included in each experiment.

### Quantitative real-time RT-PCR

Total RNA from the cultures was extracted using the Direct-zol RNA MiniPrep Kit (Zymo Research), and 0.5–1 μg RNA was reverse transcribed using the iScript cDNA Synthesis Kit (Bio-Rad). Specific primers, including the GAPDH control, were designed using IDT SciTools Real-Time PCR software. We performed gene-specific mRNA analyses using the Bio-Rad CFX96 system. Quantitative PCR amplification reactions contained the same amount of reverse transcription product, including 7.5 μl of 2× iQSYBR Green Mix (Bio-Rad) and 100–300 nM forward and reverse primers in a final volume of 15 μl. The primer sequences are listed below. Primer efficiency was obtained from the standard curve and integrated for the calculation of relative gene expression, which was based on real-time PCR threshold values of different transcripts. The primer sequences (5’ to 3’) are as follows:

*Il-1β*: forward - TTGTGGCTGTGGAGAAGCTGT, reverse - AACGTCACACACCAGCAGGTT;

*Tnf-α*: forward - AGAAGTTCCCAAATGGCCTCCCT, reverse - TACAACCCATCGGCTGGCACCAC;

*Il-10*: forward - TGTCAAATTCATTCATGGCCT, reverse - ATCGATTTCTCCCCTGTGAA;

*Gapdh* : forward - AGGTCGGTGTGAACGGATTTG, reverse - GGGGTCGTTGATGGCAACA.

### Bulk RNA sequencing and pathway analysis of DRG tissues

Total RNA was extracted from L3-L5 DRG of mice with sham surgery and fracture surgery treated with vehicle or PDX mice using RNeasy® Mini Kit (QIAGEN, Cat# 74104). RNA quantity and quality were assessed with a NanoDrop™ One spectrophotometer (Thermo Fisher Scientific). All samples displayed a 260:280 ratio greater than 2.0 and RNA Integrity Numbers (RINs) above 8.0, with RNA concentration > 150 ng/μL and the total RNA yield > 3 μg/sample. The Poly(A) RNA sequencing library was prepared according to Illumina’s TruSeq-stranded-mRNA sample preparation protocol. RNA integrity was verified using the Agilent Technologies 2100 Bioanalyzer. Poly(A) tail-containing mRNAs were purified using oligo-(dT) magnetic beads, undergoing two rounds of purification. After purification, the poly(A) RNA was fragmented in a divalent cation buffer at elevated temperature, and the DNA library was then constructed. Quality control and quantification of the sequencing library were conducted using the Agilent Technologies 2100 Bioanalyzer High Sensitivity DNA Chip. Paired-end sequencing was carried out on the Illumina NovaSeq 6000 sequencing system by LC Sciences.

RNA-seq differential gene expression data were analyzed using QIAGEN’s Ingenuity Pathway Analysis (IPA, QIAGEN Redwood City, www.qiagen.com/ingenuity) software version 01-16. Top Canonical Pathway analysis was conducted to compare PDX and vehicle treated groups with fracture. The significance of the association between the dataset and canonical pathways was measured in two ways: 1) A ratio of the number of molecules from the dataset that map to the pathway, divided by the total number of molecules that map to the canonical pathway. 2) Fisher’s exact test was used to calculate a p-value determining the probability that the association between the genes in the dataset and the canonical pathway is explained by chance alone. Volcano Plot: The volcano plot visualizes the relationship between the statistical significance (-log10 p-value) and the magnitude of change (log2 fold change) for genes in the study.

### *In vivo* recordings of C-fiber reflex using electromyogram

As previously described, an electromyogram (EMG) was used to record the C-fiber reflexes from the biceps femoris (74). Briefly, mice were anesthetized with 1.5% isoflurane, and needle electrodes were inserted subcutaneously into the medial part of the 3rd and 4th toes. The electrodes delivered electrical stimuli with single square waves of 1 ms from a constant current isolated stimulator (DS3, Digitimer Ltd, London, England) to evoke C-fiber reflexes. EMG signals were recorded using a pair of platinum-iridium electrodes inserted into the left biceps femoris muscle. These signals were amplified via a microelectrode amplifier (A-M system, Sequim, WA), and they were then recorded using an acquisition system (Digidata 1440, Molecular Devices). The C-fiber reflex threshold (Tc) was identified when the EMG signals corresponding to C-fiber activities were elicited. After detecting Tc, the EMG reflex resulting from stimulation at 2-fold Tc was established as the noxious stimulation-induced responses. C-fiber reflexes were recorded before and after 30 min PDX treatment (peri-sciatic nerve injection, 20 ng in 20 μL; intraperitoneal injection, 150 ng in 100 μL), with PBS as vehicle control.

### Computer simulations

The protein sequence of human GPR37 (hGPR37) was downloaded from the UniProt database (ID: O15354) in fasta format. Homology modeling was performed using the GPCR-ModSim server, which employs an automated approach for template selection and model generation (75). To elucidate the binding mode of all ligands in the binding site of the homology model of hGPR37, docking studies were performed with the help of Autodock4 software (76). Before the docking, the hGPR37 structure was prepared using the AutoDock Tools 4 software, which involved adding hydrogen atoms, assigning Gasteiger charges, and defining rotatable bonds. Based on previous mutagenesis studies and conservation analysis, the docking grid was centered on the putative binding site, encompassing residues identified as crucial for ligand interaction. For each ligand, 20 docking runs were performed, and the top-scoring poses were selected for further analysis. Molecular dynamics simulation and docking grid analysis were performed as previously described (19). The 100 ns of molecular dynamics simulation (MDS) was performed on the cluster 1 structure to assess the stability of the docking complex.

### Plasmids and AAV production

pAdDeltaF6 was a gift from James M. Wilson (Addgene plasmid #112867). pUCmini-iCAP-AAV.MaCPNS2 was a gift from Viviana Gradinaru (Addgene plasmid # 185137). pAAV.CAG.GCaMP6f.WPRE.SV40 (Addgene plasmid #100836) and pAAV.Syn.GCaMP6f.WPRE.SV40 (Addgene plasmid # 100837) were gifts from Douglas Kim & GENIE Project. HEK293T cells were cultured in DMEM supplemented with 10% fetal bovine and 1.5 x10^7^ cells were seeded per 15 cm dish 24 h before transfection. To produce each virus, a total of six dishes were prepared. Cells were then transfected with 30 µg pAd-DELTA F6, 15 µg serotype plasmid AAV.MaCPNS2, and 15 µg AAV plasmid with PEI MAX (Polysciences, Cat#24765). After 72 h, cells were harvested and lysed in 4 ml of lysis buffer (15 mM NaCl, 5 mM Tris-HCl, pH 8.5) using three freeze-thaw cycles, and incubated with 50 U/ml Benzonase (Millipore, Cat#70664) for 30 min at 37°C. After spinning down at 4500 rpm for 30 min at 4°C, the supernatant was added to the top of 15%, 25%, 40% and 60% stacked iodixanol gradients and centrifuged using a Beckman Ti-70 rotor, spun at 67,000 rpm for 1.5 h at 18°C. The viral solution was then collected from the interface between 40%-60% iodixanol gradients, washed with 1X PBS (Gibco, Cat#14190144), and concentrated with a 100 kDa filter (Millipore, Cat#UFC910008). The purified virus was aliquoted and stored at −80°C until use.

### Paw edema assessment and Evans blue test

The weight of mouse hind paw was used to determine paw swelling (edema) after capsaicin and AITC injection. To assess vascular permeability, which is associated with paw swelling, Evans blue dye (1%, 200 μl) was injected into the tail veins of all mice 15 minutes after the capsaicin and AITC injections (77). Photographs of the hind paws were taken 15 minutes post-injection. The paws were then placed in a 65°C oven for 48 hours to remove moisture content and ensure consistent weight measurements. Subsequently, the dried paw tissues were weighed, crushed, and soaked in 500 μl of formamide. The samples were heated in a water bath at 55°C for 24–48 hours to extract the Evans blue dye. The formamide/Evans blue mixture was centrifuged at 4000 × g for 10 minutes, and the Evans blue dye concentration in the supernatant was quantified by measuring absorbance at 630 nm using a microplate reader.

### Flow Cytometry Analysis of Efferocytosis in Bone Fracture Tissue

Wild-type (WT) and *Gpr37* knockout (KO) mice were subjected to bone fracture, and muscle tissue surrounding the fracture site was harvested 3 days post-injury. Tissues were minced and enzymatically digested in 1 mg/mL collagenase/Dispase (Roche) for 60 minutes at 37°C with gentle agitation. Following digestion, samples were filtered through a 50 μm cell strainer to obtain single-cell suspensions. Enzymatic activity was quenched using 10% fetal bovine serum (FBS), and cells were washed with PBS containing 10 mM EDTA. Cells were then resuspended and incubated for 10 minutes with Invitrogen’s permeabilization solution to allow intracellular staining. To block Fc receptors, cells were incubated with anti-CD16/32 antibody (1 µg/ml) in 10% bovine serum albumin (BSA) for 1 hour at 4°C. Cells were then stained with the following antibodies diluted 1:200 in cold PBS + 10% BSA + 10 mM EDTA: F4/80-FITC(Biolegend: 1:200) for macrophage identification, Ly6G-APC-Cy7(Biolegend, 1:200) for neutrophil detection, Hoechst dye **(Sigma)** for live/dead cell discrimination, and nuclear staining. After staining, cells were washed in PBS with EDTA. The flow cytometry events were acquired in a BD FACS Canto II flow cytometer by using BD FACS Diva 8 software (BD Bioscience). Data were analyzed using Cytobank Software (https://www.cytobank.org/cytobank). The gating strategy was illustrated in Supplemental Figure 14.

### Spatial transcriptomic analysis in human DRG

Human dorsal root ganglia (DRG) were obtained from two donors (a 63-year-old male and a 63-year-old female) through the National Disease Research Interchange (NDRI) with exemption permission from the Duke University Institutional Review Board (IRB). Postmortem L3–L4 DRG were delivered in ice-cold cell culture medium within 28 hours of death. Upon receipt, tissues were immediately dissected and fixed in 4% paraformaldehyde followed by cryoprotection in 30% sucrose at 4°C for a minimum of three nights.

Visium HD Experiment. OCT-embedded frozen DRG tissues were cryosectioned at 20 μm thickness and stored at –80°C. Slides were rehydrated, H&E stained, and destained following the 10x Genomics Visium CytAssist Spatial Gene Expression for Fixed Frozen – Rehydration, H&E Staining, Imaging & Decrosslinking protocol (CG000662). Brightfield imaging was performed using a Zeiss Axioscan Z1 slide scanner with a 20× objective. Following imaging, tissue sections were decrosslinked according to the *Visium HD FFPE Tissue Preparation Handbook* (CG000684) and then processed using the *Visium HD Spatial Gene Expression Reagent Kits* (CG000686).

In brief, whole transcriptome probe panels were applied to each section for hybridization and ligation across the transcriptome. Slides were loaded into the Visium CytAssist instrument, which enabled alignment of the tissue section with the Visium HD slide. Gene expression probes and antibody tags were released through CytAssist-enabled RNA digestion and tissue removal, allowing capture by spatially barcoded oligonucleotides on the slide surface. The slides were then removed for extension and downstream library preparation. Barcoded ligation products were amplified, purified, and indexed by PCR for multiplexing, with the optimal number of cycles determined by qPCR. Final libraries were quantified using KAPA qPCR and sequenced on an Illumina NovaSeq X Plus 1.5B flow cell with the following read lengths: Read 1: 43 cycles; i7 Index: 10 cycles; i5 Index: 10 cycles; Read 2: 50 cycles.

Data Processing and Analysis: Raw FASTQ files were processed using Space Ranger software (v3.1.2, 10x Genomics) for alignment to the reference transcriptome and generation of gene expression matrices at 2 μm, 8 μm, and 16 μm binning resolutions. CytAssist images were manually aligned to high-resolution H&E images using the Loupe Browser (10x Genomics) prior to running the spaceranger count command. Inputs included the CytAssist image, microscope image, FASTQ files, probe set, reference transcriptome, and alignment JSON file. Outputs included spatially resolved gene expression matrices and annotated tissue images.

